# Neuron Type-Specific Translatomes in Spinal Cords of Naïve and Neuropathic Mice

**DOI:** 10.1101/2020.03.19.996645

**Authors:** R.R. Das Gupta, L. Scheurer, P. Pelczar, W.T. Ralvenius, H. Wildner, H.U. Zeilhofer

## Abstract

The spinal dorsal horn harbors a sophisticated and heterogeneous network of excitatory and inhibitory neurons that process peripheral signals encoding different sensory modalities. Although it has long been recognized that this network is crucial both for the separation and the integration of sensory signals of different modalities, the molecular identity of the underlying neurons and signaling mechanisms are still only partially understood. Here, we have used the translating ribosome affinity purification (TRAP) technique to map the translatomes of excitatory glutamatergic (VGLUT2^+^) and inhibitory GABA and/or glycinergic (VGAT^+^ or Gad67^+^) neurons of the mouse spinal cord. Our analyses demonstrate that inhibitory and excitatory neurons are primarily set apart by the expression of genes encoding transcription factors or genes related to the production, release or re-uptake of their principal neurotransmitters (glutamate, GABA or glycine). Subsequent gene ontology (GO) term analyses revealed that neuropeptide signaling-related GO terms were highly enriched in the excitatory population. Eleven neuropeptide genes displayed largely non-overlapping expression patterns closely adhering to the laminar and hence also functional organization of the spinal cord grey matter, suggesting that they may serve as major determinants of modality-specific processing. Since this modality-specific processing of sensory input is severely compromised in chronic, especially neuropathic, pain, we also investigated whether peripheral nerve damage changes the neuron typespecific translatome. In summary, our results suggest that neuropeptides contribute to modality-specific sensory processing in the spinal cord but also indicate that altered sensory encoding in neuropathic pain states occurs independent of major translatome changes in the spinal neurons.

## Introduction

The ability to sense and discriminate different noxious and innocuous somatosensory stimuli is essential for all higher animals and humans in order to react adequately to external stimuli and internal conditions [1, 2]. The spinal dorsal horn, i.e., the sensory part of the spinal cord, constitutes a key element in this process. It receives somatosensory signals from peripheral neurons and processes these signals together with other inputs descending from supraspinal sites in a complex network of inhibitory and excitatory interneurons before relaying these signals via projection neurons to supraspinal centers [3]. Projection neurons make up less than 10% of all dorsal horn neurons, while more than 90% of the neuronal population are interneurons of which, between 60 and 70% are excitatory glutamatergic neurons, and the rest is inhibitory (GABA and/or glycinergic).

The spinal cord is organized in a laminar fashion, which has initially been proposed on the basis of differences in cell density and morphology between the different laminae [4, 5] but has later been shown to also reflect functional organization. This is especially reflected for example by the lamina specific innervation pattern by the different types of peripheral sensory neurons: unmyelinated C fibers, which mainly carry noxious and thermal information, terminate in the superficial dorsal horn (laminae I-II), while thickly myelinated Ab fibers, which convey innocuous signals including touch and proprioceptive information, terminate in the deep dorsal horn (laminae III-V) [3, 6, 7]. A laminar organization of neuronal function is also supported by gene expression patterns that follow laminar patterns [8–12]. Furthermore, optogenetic and chemogenetic experiments support a modality-specific processing by distinct genetically defined neuron populations [7, 13–15]. However, the cellular basis of this modality specific processing and hence the identity of interneuron types is only incompletely understood. Recent work has used single cell RNA sequencing and unsupervised clustering to identify 15 subtypes or excitatory and 15 types of inhibitory neurons [8]. In the present study, we employed translating ribosome affinity purification (TRAP) technology [16] to characterize the translatomes of excitatory and inhibitory spinal neurons. To this end, we generated VGLUT2::bacTRAP (*Slc17a6*), VGAT::bacTRAP (*Slc32a1*) and Gad67::bacTRAP (*GAD1*) mice, which express the eGFP-tagged ribosomal subunit L10a (RPL10a) under the control of the respective gene regulatory elements. Comparing the translatomes of excitatory and inhibitory spinal neurons by differential gene expression analysis in VGLUT2::bacTRAP and Gad67::bacTRAP mice revealed two classes of genes that distinguish excitatory and inhibitory neurons, namely transcription factors and genes related to the neurotransmitter phenotype of these neurons. In addition, we found that genes encoding neuropeptides constitute a functionally defined class of genes that is highly enriched in excitatory spinal neurons. Accordingly, multiplex *in situ* hybridization for neuropeptide encoding genes revealed largely non-overlapping expression patterns that follow the laminar organization of the spinal cord thus suggesting a role of neuropeptide signaling in segregation of sensory modalities.

Interestingly, under pathophysiological conditions, such as neuropathy, the segregation of innocuous and noxious information becomes compromised. Excitatory interneurons of the deeper dorsal horn engage in circuits that process noxious information and thereby contribute to the development of mechanical hypersensitivity or allodynia (i.e. pain induced by innocuous sensory stimuli) [17–20]. Several lines of evidence support the idea that these circuit changes are based on the loss of inhibition exerted by e.g. glycinergic and/or parvalbumin expressing inhibitory interneurons [19, 21]. The underlying molecular mechanisms inducing these circuit changes are poorly understood. We have therefore employed the TRAP technology to search for molecular changes that occur in neuropathic mice. To this end, we compared the translatomes of VGLUT2::bacTRAP and Gad67::bacTRAP mice isolated from mice 7 days after sham surgery or mice 7 days after inducing a neuropathy either by the chronic constriction injury (CCI) or spared nerve injury (SNI). While we found hundreds of genes that distinguish inhibitory and excitatory spinal neurons, our results failed to reveal consistent translatome changes in spinal dorsal horn neurons in two well-established mouse neuropathy models. They thus suggest that neuropathy-induced changes in dorsal spinal sensory circuits occur independent of changes in the translatomes of dorsal horn neurons.

## Results

### Generation and validation of three bacTRAP mouse lines to profile gene translation in excitatory and inhibitory neurons

In this study, we set out to determine the translatome (polysome-bound mRNA) of excitatory and inhibitory neurons of the spinal cord. To this end, we generated three bacTRAP mouse lines; VGLUT2::bacTRAP (Tg(Slc17a6-RPL10a-eGFP)Uze), Gad67::bacTRAP (Tg(Gad1-RPL10a-eGFP)Uze) and VGAT::bacTRAP mice (Tg(Slc32a1-RPL10a-eGFP)Uze). In these mice, the eGFP-tagged ribosomal protein L10a is expressed as a transgene either in glutamatergic neurons (VGLUT2::bacTRAP (Fig 1A, D, G), all inhibitory neurons (GABAergic and glycinergic neurons) (VGAT::bacTRAP, Fig 1B, E, H) or in GABAergic neurons that utilize Gad67 for GABA synthesis (Gad67::bacTRAP, Fig 1C, F, I).

**Figure 1.**
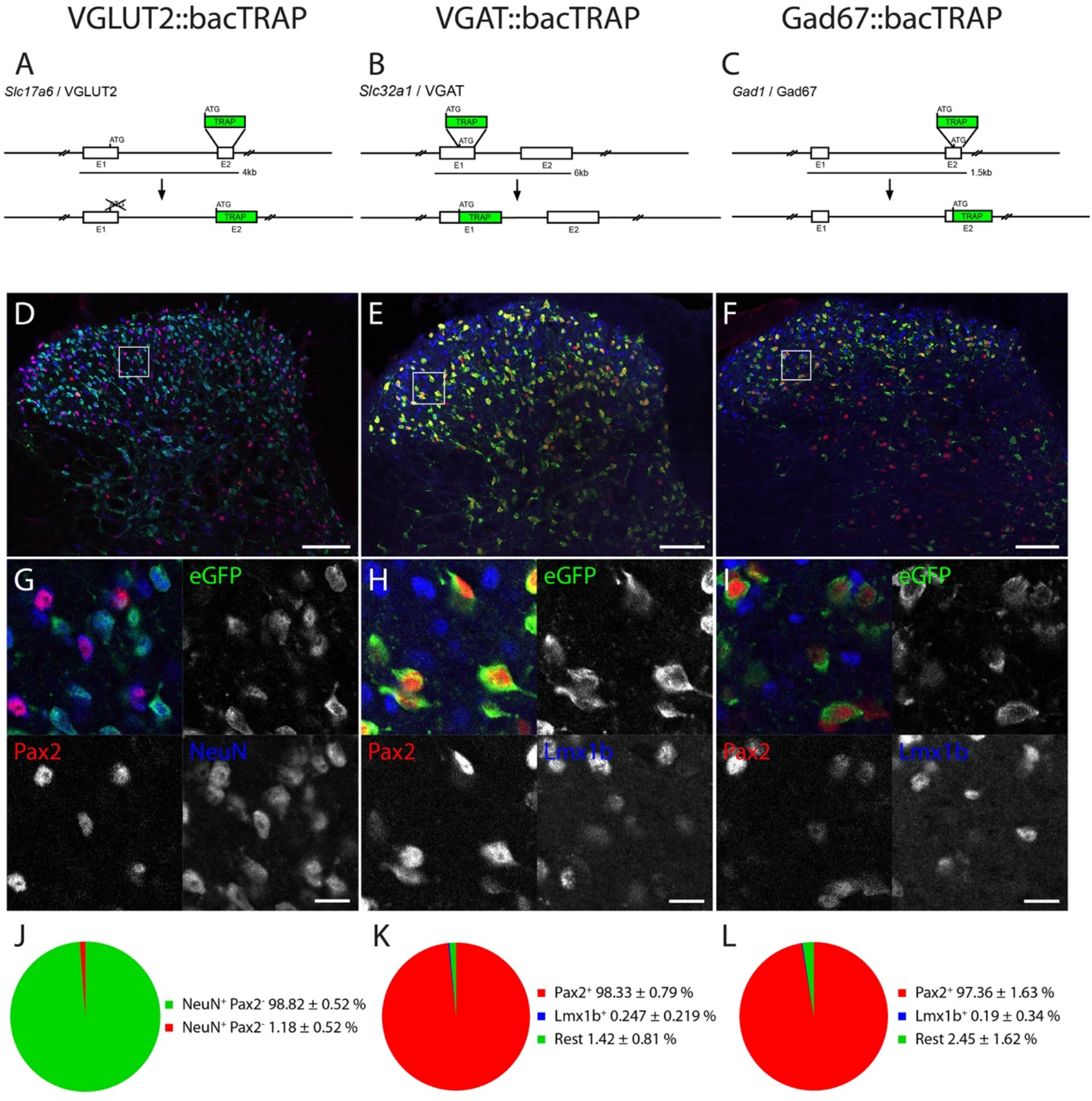
Generation and validation of three bacTRAP mouse lines, specific to excitatory and inhibitory spinal cord neurons. (A, D, G, J) VGLUT2::bacTRAP line, (B, E, H, K) VGAT::bacTRAP line, (C, F, I, L) Gad67::bacTRAP line. (AC) BAC constructs for the three mouse lines. (D-I) Immunofluorescence images of the lumbar spinal dorsal horn of the three mouse lines. Boxes in the overview images of (D-F) indicate the position of the higher magnification images in (G-H). (D, G) VGLUT2::bacTRAP line with eGFP in green, Pax2 in red and NeuN in blue. (E, H) VGAT::bacTRAP mouse line with eGFP in green, Pax2 in red and Lmx1b in blue. (F, I) Gad67::bacTRAP mouse line with eGFP in green, Pax2 in red and Lmx1b in blue. (J) Quantification of eGFP+ NeuN+ cells with and without signal for Pax2 in the VGLUT2::bacTRAP line. (K-L) Quantification of eGFP+ cells with signal for Pax2 or Lmx1b in (K) the VGAT::bacTRAP line and (L) the Gad67::bacTRAP line. (D-G) Scale bar = 100 μm. (G-H) Scale bar = 5 μm.

From the spinal cord of these mice, polysome-bound mRNA can be isolated by coimmunoprecipitation of mRNA bound to the eGFP-L10a tagged ribosomes. We confirmed the correct expression of the eGFP-L10a transgene in the lumbar spinal dorsal horn of the different mouse strains by immunohistochemistry. NeuN was used as a neuronal marker, Pax2 as a ubiquitous marker for spinal inhibitory [21, 22] and Lmx1b as a marker for the majority of dorsal spinal excitatory neurons (Fig 1D-I). In the VGLUT2::bacTRAP line, the eGFP-L10a expression is restricted to Pax2-negative neurons (98.8 ± 0.5% Pax2^-^ NeuN^+^; Fig 1D, G, J), indicating its exclusive expression in excitatory spinal neurons. In the two mouse lines targeting inhibitory neurons, VGAT::bacTRAP and Gad67::bacTRAP, the expression was confined to Pax2-positive neurons (98.3 ± 0.8%, 97.4 ± 1.6%, respectively) and virtually absent from excitatory, Lmx1b-positive neurons of the dorsal horn (0.25 ± 0.22%, 0.19 ± 0.34%, for VGAT::bacTRAP and Gad67::bacTRAP mice, respectively; Fig 1E-F, H-I, K-L). These expression patterns resemble those described in *in situ* hybridization experiments (Allen Brain Atlas, mousespinal.brain-map.org) with widespread expression throughout the entire dorsal horn in the VGLUT2::bacTRAP line and the VGAT::bacTRAP line, and more localized expression in the superficial dorsal horn in Gad67::bacTRAP mice.

### General differences in gene expression in excitatory and inhibitory spinal neurons

We used the bacTRAP mouse lines to search for genes that could potentially serve as markers for subpopulations of excitatory or inhibitory dorsal horn neurons. To this end, we isolated and sequenced the cell-type-specific polysomal mRNA from the lumbar spinal cord of three animals from each mouse line. Out of 22’880 detectable genes, between 13’515 and 13’774 were found expressed in each mouse line (Fig 2A, Table S1), defined by a normalized count (number of reads per gene, normalized to the total number of reads and the gene length) of > 10. In order to identify among the expressed genes those that show a regionally non-overlapping expression pattern, we focused our analyses on genes whose regional expression is accessible by *in situ* hybridization. To this end, we assumed that genes with normalized counts > 100 should be reliably detectable with *in situ* hybridization. This threshold is three times lower than the expression of *Grpr*, a gene that we previously found expressed at low levels in a small subset of dorsal horn neurons. In all three mouse lines, approximately 11’200 genes were expressed at levels exceeding this threshold (Fig 2A).

**Figure 2.**
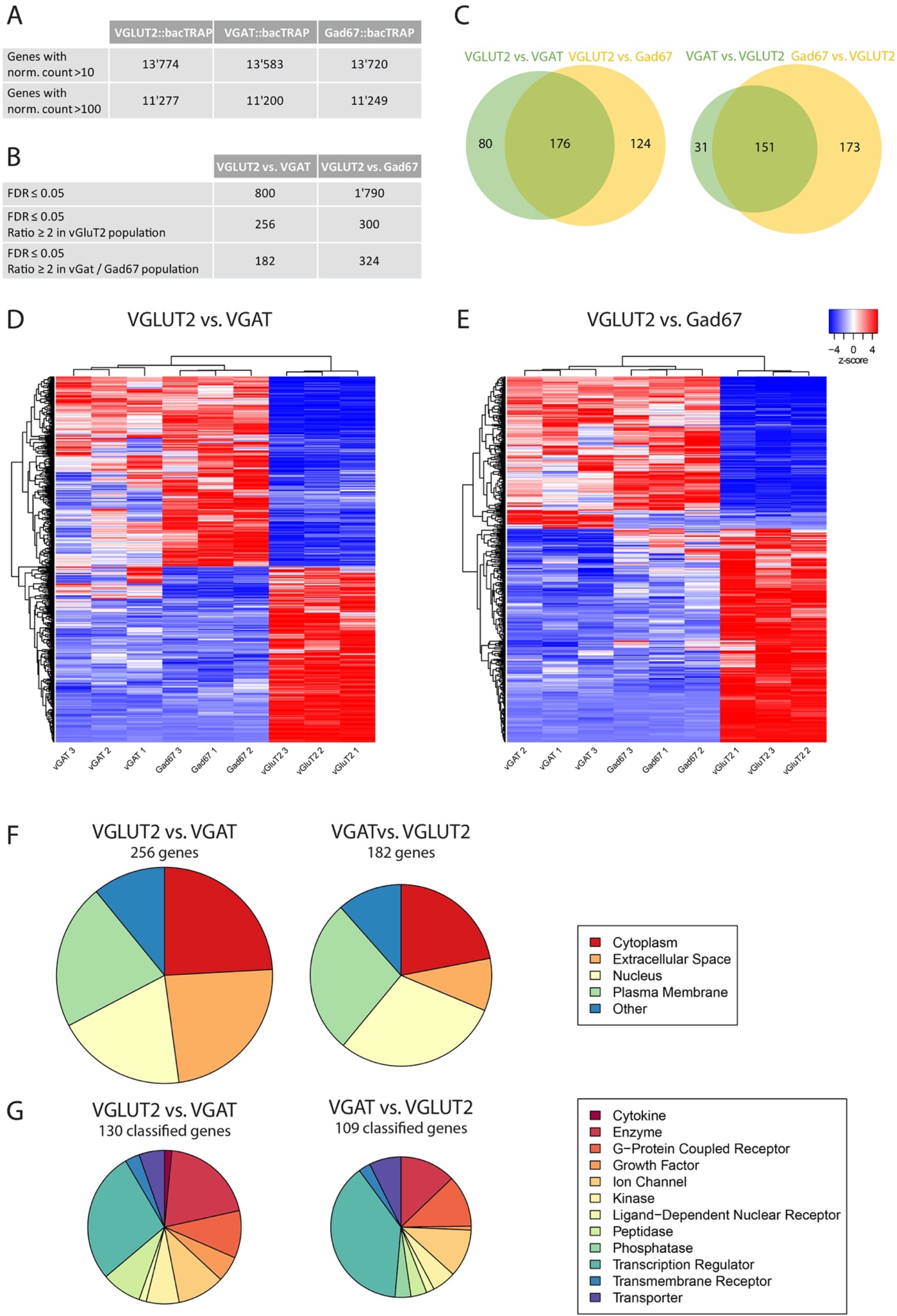
RNA-Seq metadata from sequencing of the polysomal mRNA from VGLUT2::bacTRAP, VGAT::bacTRAP and Gad67::BacTRAP mice. (A) Number of “expressed” genes (> 10 norm. counts) and number of genes with > 100 norm. counts in the three different mouse lines. (B) Number of significantly enriched genes (FDR ≤ 0.05) in the DGEAs between VGLUT2::bacTRAP and VGAT::bacTRAP / Gad67::bacTRAP. Number of significantly enriched genes with ratio ≥ 2 in the excitatory and inhibitory neurons of both DGEAs. (C) Congruence between the enriched genes of the two DGEAs: 176 genes in common for the genes enriched in VGLUT2 neurons and 151 genes in common for the genes enriched in the inhibitory neurons. (D-E) Cluster analysis with the nine mRNA samples and the genes differentially expressed in (D) VGLUT2 vs. VGAT (800 genes, FDR ≤ 0.05) and (E) VGLUT2 vs. Gad67 (1790 genes, FDR ≤ 0.05). Color key indicated z-score with white representing zero, blue representing negative z-scores and red representing positive z-scores. (F) Subcellular location and (G) coarse function of the genes enriched in VGLUT2 vs. VGAT and VGAT vs. VGLUT2. Genes with unclassified function are not displayed.

Differential gene expression analyses (DGEAs) revealed 800 genes that were differentially expressed (with a false discovery rate (FDR) ≤ 0.05) in VGLUT2::bacTRAP versus VGAT::bacTRAP mice, and 1’790 genes in VGLUT2::bacTRAP versus Gad67::bacTRAP mice (Fig 2B). The approximately two-fold lower number of significantly enriched genes in the comparison of VGLUT2::bacTRAP versus VGAT::bacTRAP mice probably results from the lower expression of the eGFP-L10a transgene in the VGAT::bacTRAP line and hence a lower mRNA yield and higher variability in read numbers.

A comparison of the translatomes of VGLUT2 positive and VGAT positive neurons revealed 438 differentially expressed genes with an at least two-fold enrichment in one of the two populations. Of those, 256 genes were enriched in excitatory neurons, and 182 in inhibitory neurons. A higher number of differentially expressed genes (300 and 324) were detected when the Gad67-TRAP line was used instead of VGAT-TRAP line (Fig 2B). Comparing the two DGEAs revealed that about half of the genes enriched in either the excitatory or the inhibitory populations and identified in the comparison with VGAT::bacTRAP line were also found when the Gad67::bacTRAP line was used (Fig 2C) supporting the robustness of our approach.

Next, we performed cluster analyses ([23] (http://fgcz-shiny.uzh.ch/fgcz_heatmap_app/)) of the 800 genes differentially expressed in VGLUT2::bacTRAP and VGAT::bacTRAP mice (Fig 2D), and of the 1790 genes differentially expressed in VGLUT2::bacTRAP and Gad67::bacTRAP mice (Fig 2E) over the nine different samples. Clustering of the samples was similar for both comparisons. As expected, the first hierarchical segregation separated the inhibitory (VGAT and Gad67) from the excitatory (VGLUT2) samples. Within the inhibitory cluster VGAT samples segregated from Gad67 samples (Fig 2D-E). The cluster analysis therefore identified the VGAT^+^, Gad67^+^ and VGLUT2^+^ samples as three populations with distinct translatome profiles.

### Transcription factor expression and neuropeptide expression are key features distinguishing excitatory and inhibitory spinal neurons

To identify key features that distinguish excitatory and inhibitory neurons beyond the expression differences of individual genes, we gathered information on the subcellular location and function of the enriched genes as well as overrepresented pathways (Fig 2 F-G). To this end, we first employed the Ingenuity Pathway Analysis (IPA, QIAGEN) tool, which assigns a unique subcellular location (“location”) and coarse function (“type(s)”) to the enriched genes. Twenty-four percent of the genes enriched in VGLUT2 over VGAT encoded for cytoplasmic proteins, 24% for extracellular proteins, 22% for proteins in the plasma membrane, 20% were nuclear proteins and the remaining 11% were located elsewhere. Similar localizations were found when genes enriched in the inhibitory populations were analyzed. One hundred-thirty out of 256 genes enriched in VGLUT2 over VGAT and 109 of 182 genes enriched in VGAT over VGLUT2 could be assigned to a coarse function (Fig. 2G). The largest group of genes that distinguished inhibitory from excitatory neurons in the adult spinal cord were transcription regulators (28% and 39% of the genes enriched in VGAT and VGLUT2, respectively). Many of them (such as *Tlx3*, *Lmx1b*, *Phox2a* and *Prrxl1* expressed in excitatory or *Lhx1/5*, *Pax2* and *Gbx1* expressed in inhibitory neurons) have been described as crucial regulators of cell fate determination of spinal neurons during development [15, 24–31]. In addition, we found some transcription factors with as yet unknown functions in spinal neurons such as *Tfap2b* and *Sall3*. Two other large molecular groups present in the set of enriched genes were enzymes and G protein coupled receptors.

A subsequent Gene Ontology (GO) term overrepresentation analysis for the molecular function [32, 33] confirmed an enrichment of molecules involved in transcription and neuropeptide signaling. In all four comparisons, several molecular function terms related to the regulation of transcription were overrepresented, e.g. *proximal promoter DNA-binding, transcription activator activity, RNA polymerase II-specific* and *sequence-specific DNA binding*, *E-box binding* (Fig 3 A-D). Another set of highly overrepresented molecular function terms related to neuropeptide signaling (e.g. *neuropeptide receptor activity* and *neuropeptide receptor binding)*. In all comparisons, neuropeptide signaling-related molecular functions terms were highly overrepresented (4.6 – 26.3-fold overrepresented as compared to all expressed genes, FDR between 0.00016 and 0.0488), and those related to transcription regulation included the largest numbers of enriched genes (1.9 – 10.3-fold overrepresented, FDR 0.0000873 – 0.0445). Among the 50 most enriched genes (Table S2–5) filtered for genes with a normalized count of ≥ 100 in at least two out of three samples of any mouse line, transcription factors constituted the most abundant molecular category (30 – 38% for the different comparisons), followed by genes that encode neuropeptides (24 – 26%).

**Figure 3.**
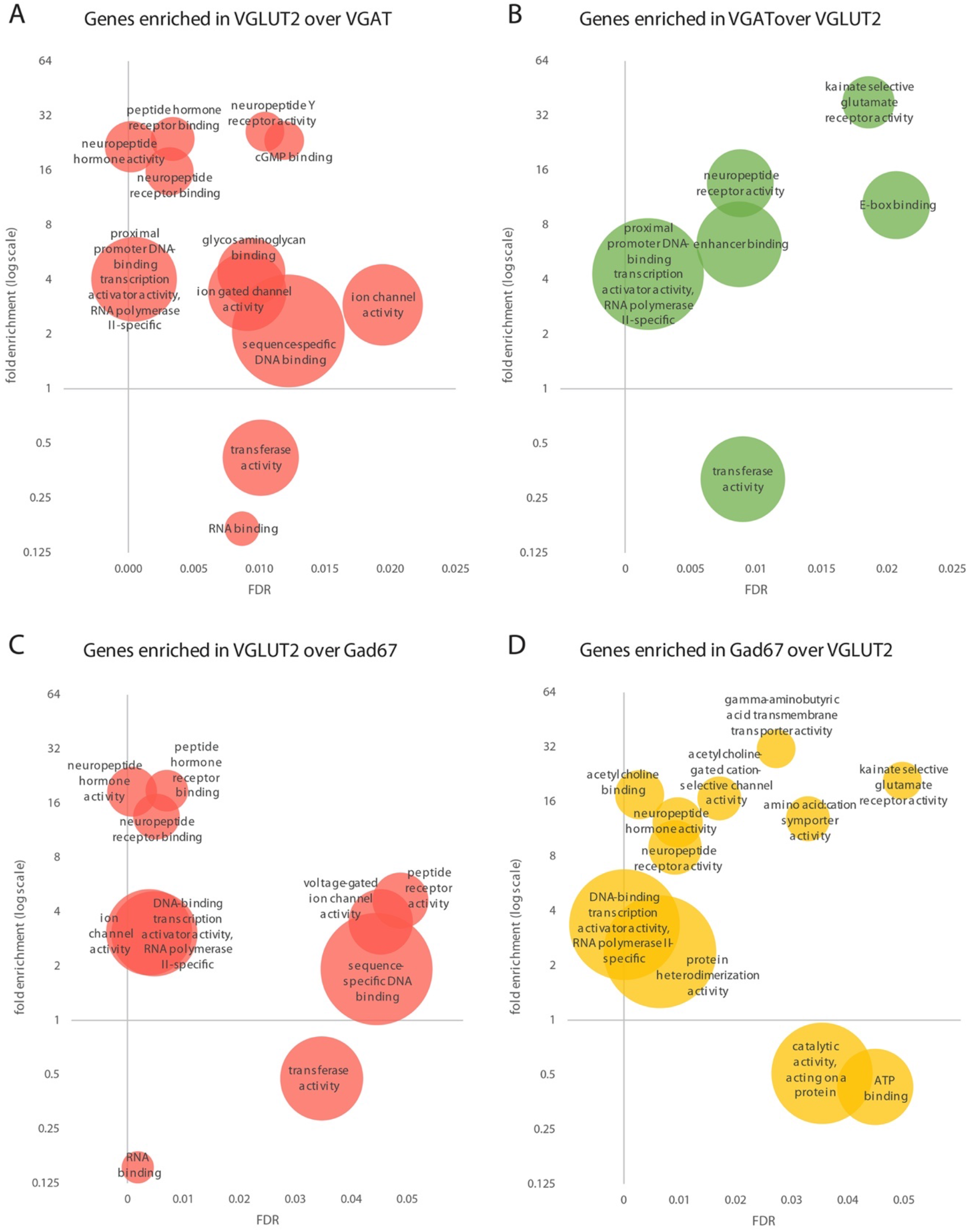
GO term overrepresentation analyses for the molecular function (MF) with the enriched genes in the comparison of VGLUT2::bacTRAP to VGAT::bacTRAP and Gad67::bacTRAP. Overrepresented MF GO terms in the genes enriched in the comparison of (A) VGLUT2 vs. VGAT, (B) VGAT vs. VGLUT2, (C) VGLUT2 vs. Gad67 and (D) Gad67 vs. VGLUT2. (A-D) x axis displays false discovery rate (FDR) of the overrepresented GO term, y axis displays fold overrepresentation on a logarithmic scale and circle size indicates number of enriched genes. When several GO terms of a hierarchy are significantly overrepresented, only the most specific GO term is displayed. (D) For comprehensibility two MF are not displayed: “acetylcholine receptor activity”, 13.12-fold enrichment, FDR = 0.00801, no of enriched genes = 5, and “RNA polymerase II proximal promoter sequence-specific DNA binding” 2.79-fold enrichment, FDR = 0. 0.000603, no of enriched genes = 27.

### Differential use of neurotransmitter signaling pathways

In the inhibitory populations, molecular function terms associated with neurotransmitter signaling and transport were found overrepresented. *Kainate selective glutamate receptor activity* is overrepresented in both inhibitory populations (38-fold in VGAT versus VGLUT2, FDR = 0.019, 21-fold in Gad67 versus VGLUT2, FDR = 0.0497). Acetylcholine signaling related molecular function terms are overrepresented in Gad67 versus VGLUT2 (13 – 17-fold, FDR = 0.0029 – 0.0171). *Gamma-aminobutyric acid transmembrane transporter activity* is 31.5-fold overrepresented in the comparison of Gad67 to VGLUT2 (FDR = 0.027). Ion channel activity-related molecular function terms are overrepresented within the genes enriched in both comparisons of the excitatory to the inhibitory populations*. Ion gated channel activity* and *ion channel activity* in VGLUT2 versus VGAT (3.5 and 2.9-fold, FDR = 0.009 and 0.0194, respectively) as well as *voltage-gated ion channel activity* and *ion channel activity* in VGLUT2 vs. Gad67 (3.5 and 3.1-fold, FDR = 0.0452 and 0.0038, respectively).

The GO term analysis revealed an enrichment of kainate-mediated signaling and acetylcholine mediated signaling in inhibitory spinal neurons. To further complement this analysis, we performed Ingenuity Pathway Analysis on the list of the enriched genes (Fig 4). Within the 15 most significantly overrepresented pathways of each comparison additional receptor signaling pathways were found in the different neuron populations. *Glutamate* and *GABA receptor signaling* as well as *serotonin receptor signaling* were overrepresented in inhibitory neurons. As expected, the enrichment of GABA receptor signaling was mainly due to genes involved in the production (Gad65, Gad67) and transport (VGAT, Gat1) of GABA, while GABA receptors showed relatively similar expression levels in the different populations. Overrepresentation of serotonin receptor signaling (Fig. 4B) is due to the enrichment of serotonin receptors (Htr1a, Htr1d, Htr3a) in inhibitory neurons. Serotonin receptor 3a (Htr3a), the only ionotropic serotonin receptor, was highly enriched in inhibitory spinal interneurons (Table S12) and displayed an expression pattern that was restricted to a population in the deep dorsal horn (Fig S1). *GPCR signaling* as such is also overrepresented in the inhibitory enriched genes. The genes contributing to the overrepresentation encode receptors (*Grm2/3, Glp1r, Chrm3* and others) and many downstream signaling molecules (*Ptk2b, Rgs16, Plcb1, Pde1a* and others). Further enriched pathways relate to general functions of neuronal activity, such as calcium signaling (Gad67 versus VGLUT2), synaptic long-term depression (VGLUT2 versus VGAT and Gad67), axonal guidance signaling (all four comparisons) and gap junction signaling (Gad67 versus VGLUT2) (Fig 4 A-D). To further illustrate differential expression of some neurotransmitter pathways, we compiled tables indicating the relative expression levels and enrichment of glutamate, GABA, glycine, acetylcholine, serotonin, adrenergic, dopaminergic and histamine receptors (Table S5–15).

In summary, several receptor-signaling pathways are enriched within the differentially expressed genes, indicating that the respective neurotransmitter signaling pathways may engage inhibitory and excitatory spinal neurons differentially.

### Correlation of bacTRAP based translatome analysis and single cell sequencing

To relate our data to recently published single cell sequencing analysis of spinal dorsal horn neurons, we compared the 50 most enriched genes of the different comparisons to the single cell sequencing data of [8] (Table S1–4). We found that many of our hits were also present in the single cell data (60%- 70% of the 50 most enriched genes). Fourteen to twenty genes identified per comparison in our study were either not detected or only at a level that was too low to assign them to a subpopulation. Among those genes that were not detected by the single cell analysis were many transcription factors such as *Vsx2*, *Lmx1b*, *Pax5*, *Barhl1* or *Phox2a* but also some others (e.g. *Ighg3*). To verify the expression of some of these genes, we selected three for further investigation with *in situ* hybridization. We verified expression of *Barhl1* and *Phox2a*, which we found enriched in excitatory cells and the immunoglobulin heavy chain gene *Ighg3*, which we surprisingly found enriched in inhibitory neurons (Fig S2).

### A neuropeptide code for glutamatergic spinal dorsal horn neurons

We have identified transcription factors and neuropeptides as key genes differentiating inhibitory and excitatory spinal neurons. While these transcription factors are likely to impact many aspects of cellular identity, the neuropeptides probably affect communication between neurons. They therefore directly influence information processing and relay and could thus be key elements of sensory circuits. We therefore decided to more deeply investigate the expression patterns of 11 neuropeptide encoding genes identified in excitatory neurons: cholecystokinin (*Cck*), gastrin releasing peptide (*Grp*), neuromedin B (*Nmb*), neuromedin S (*Nms*), neuromedin U (*Nmu*), neuropeptide FF (*Npff*), neurotensin (*Nts*), tachykinin 1 (*Tac1*, encoding PPTA, the precursor of substance P and neurokinin A), tachykinin 2 (*Tac2*, encoding PPTB, the precursor of neurokinin B), thyrotropin releasing hormone (*Trh*) and urocortin 3 (*Ucn3*). We paid particular attention to a potential regional segregation of neuropeptide expression patterns. To this end, we conducted multiplex, fluorescent *in situ* hybridization (RNAScope), combining three different neuropeptides at a time. Figure 5 displays five of these combinations, in which every neuropeptide is represented at least once. As seen in Figure 5A, E, I, M, Q, most neuropeptides showed an expression pattern that was restricted to one or two spinal laminae. For example, *Npff*, *Ucn3*, *Nmu*, *Tac2* and *Trh* showed expression patterns largely restricted to one lamina, whereas the expression of *Grp*, *Tac1*, *Nts* and *Cck* extended over several laminae. In case of the remaining two neuropeptides, *Nmb* or *Nms*, only few cells per hemi-section exhibited high expression levels, but many cells showed weak but rather dispersed expression over most of the dorsal horn. The number of those weakly expressing cells displayed a large variability between sections. By direct comparison of the different expression patterns, we developed an approximate superficial to deep (dorsal to ventral) expression map of the neuropeptides (from superficial to deep dorsal horn): *Npff / Grp / Tac1 / Nmu / Tac2 / Ucn3 / Trh / Nts / Cck.*

**Figure 4.**
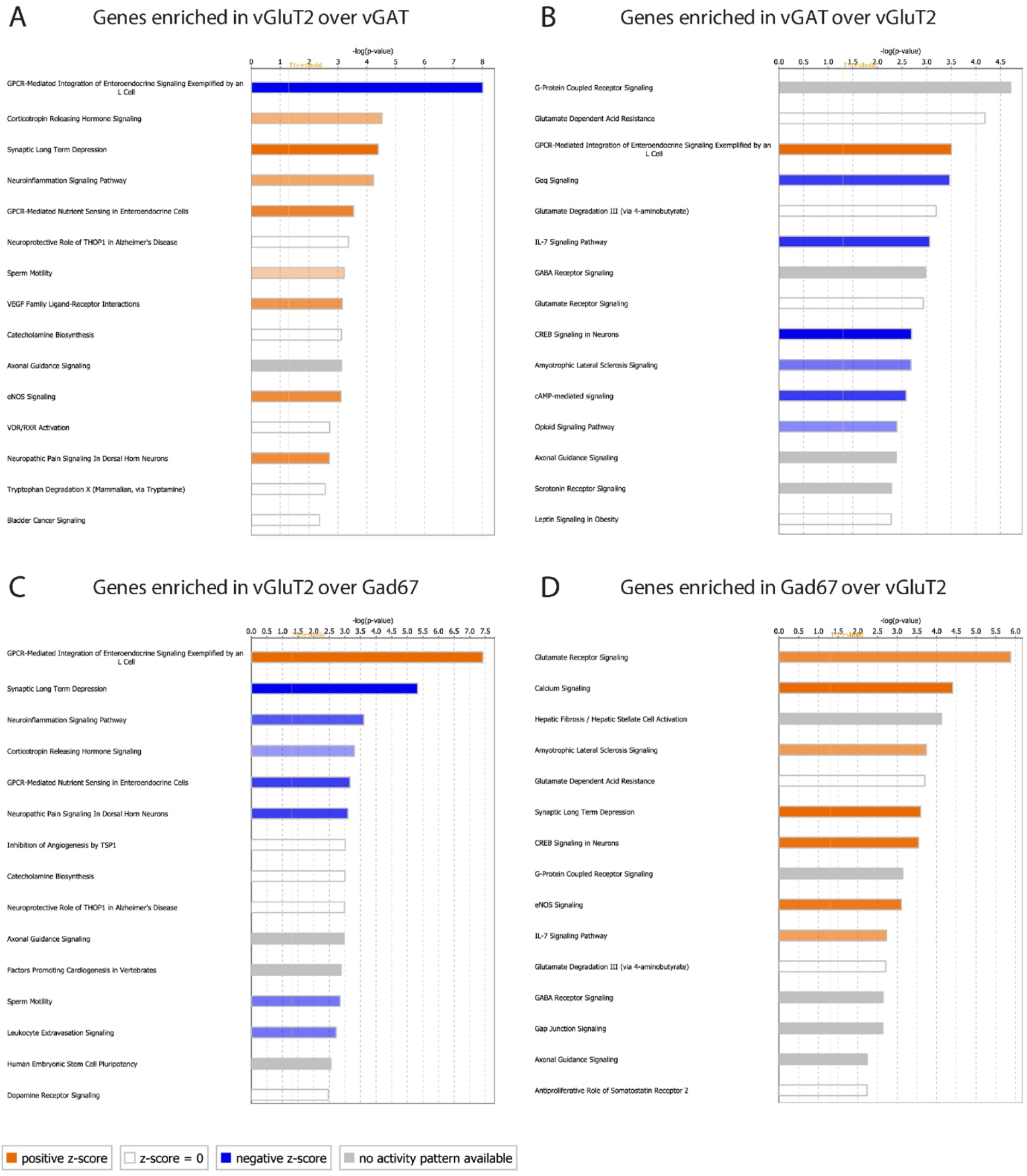
Pathway analyses with the enriched genes in the comparison of VGLUT2::bacTRAP to VGAT::bacTRAP and Gad67::bacTRAP. 15 most significantly enriched pathways within the genes enriched in the comparison of (A) VGLUT2 vs. VGAT, (B) VGAT vs. VGLUT2, (C) VGLUT2 vs. Gad67 and (D) Gad67 vs. VGLUT2. (A-D) Bar length indicates significance, color key indicates activation z-score. Orange: increased activity of the pathway predicted. Blue: decreased activity of the pathway predicted. White: z-score = zero or very close to zero (overall pathway activity is neither increased nor decreased). Gray: no prediction on pathway activity possible.

**Figure 5.**
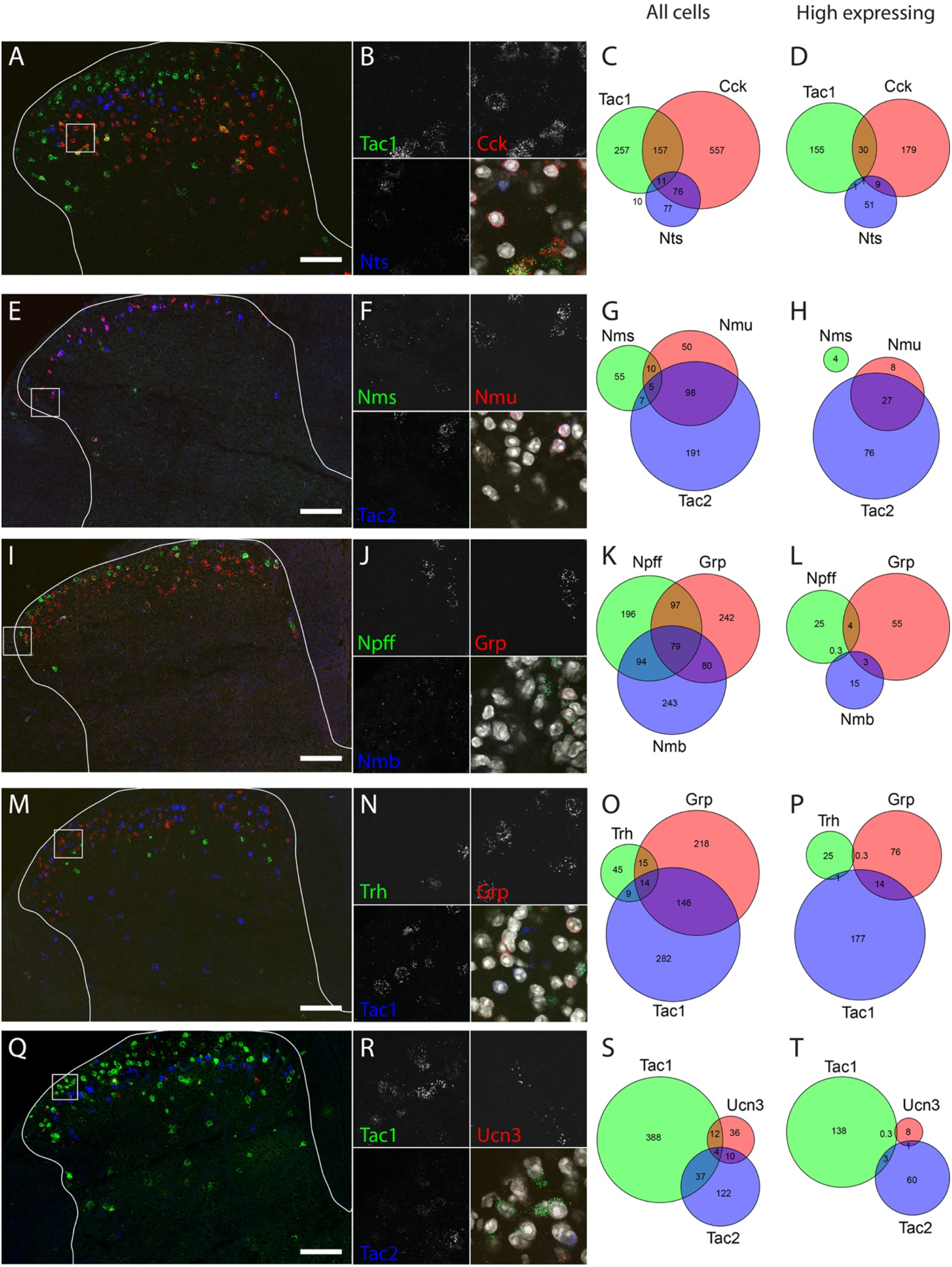
Multiplex *in situ* hybridization shows co-expression of eleven neuropeptides. Five combinations of three neuropeptides each: (A-D) Tac1, Cck and Nts; (E-H) Nms, Nmu and Tac2; (I-L) Npff, Grp and Nmb; (M-P) Trh, Grp and Tac1; (Q-T): Tac1, Ucn3 and Tac2 in green, red and blue, respectively. (A, E, I, M, Q) Overview images over the lumbar dorsal horn. The outline of the gray matter is indicated. Scale bar = 100 μm. Box indicates position of the higher magnification image in (B, F, J, N, R). (C, G, K, O, S) Number of cells per animal (two hemi-sections) that (co-)express the three neuropeptides. Circle and intersection sizes are approximately proportional to the cell number they represent. (D, H, L, P, T) Number of cells per animal that (co-)expression the three neuropeptides at high levels (> 20 dots per cell).

To determine the overlap between the different neuropeptide-expressing populations, we quantified for each pair of neuropeptides the percentage of co-expressing neurons. As some neuropeptides displayed a wide range of expression levels, we performed separate analyses either including low expressors (threshold near to background levels) or taking only cells into account with an expression clearly above background (threshold > 20 dots per cell). We created two expression matrices summarizing the percent co-expressing neurons irrespective of the neuropeptide expression level (Fig 6A) or taking only medium to high expression levels into account (Fig 6B). Restricting the analysis to the medium to high expressors led to more pronounced segregations, accentuating the absence or presence of co-expression. When focusing on medium to high-level expression, our analyses suggest that eight different subpopulations can be defined based on their neuropeptide expression pattern: 1. *Npff*^+^, 2. *Grp*^+^, 3. *Tac1*^+^, 4. *Tac2*^+^;*Nmu*^-^, 5. *Tac2*^+^;*Nmu*^+^; 6. *Nts*^+^, 7. *CCK*^+^,*Trh*^-^,*Ucn3*^-^, 8. *CCK*^+^;*Ucn3*^+^;*Trh*^+^. *Trh* and *Ucn3* display a high degree of overlap with each other (58% and 72%, respectively) and also highly overlap with *Cck* (86% and 91%, respectively). Conversely, only a minority of *Cck* neurons are *Trh* or *Ucn3* positive, suggesting that the *Trh* and *Ucn3* expressing neurons are a subpopulation of the CCK expressing neurons. Similarly, *Nmu* expressing neurons appear to define a subset of *Tac2* positive neurons (76% of the *Nmu* neurons are also *Tac2* positive and 25% of the *Tac* neurons are *Nmu* positive). In conclusion, nine out of eleven neuropeptides display a laminar expression pattern. Together, the expression of all 11 neuropeptides spans most of the dorsal horn (lamina I to V).

**Figure 6.**
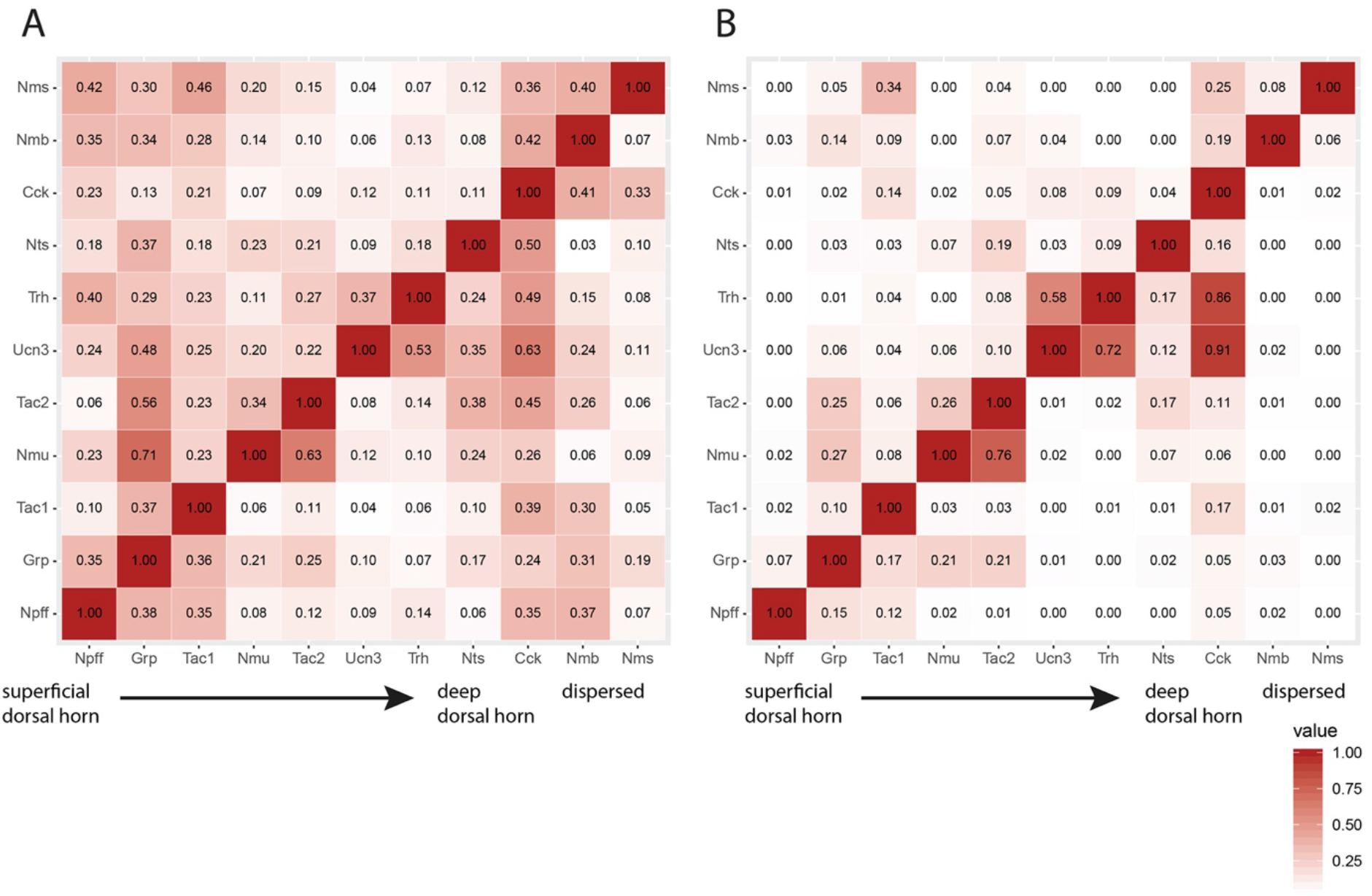
Co-expression matrices of eleven neuropeptides expressed in excitatory neurons. (A) Co-expression within all cells expressing a neuropeptide. (B) Co-expression within cells expressing clearly above background (“high levels” > 20 dots/cell) of a neuropeptide. (A-B) Fraction of cells expressing the neuropeptide on the left, that also express the neuropeptide at the bottom. For example, in (A) 42 % of all Nms-expressing cells, also express Npff, whereas in (B) 0 % of the high expressing Nms cells, also express high levels of Npff. Color key indicates co-expression with light colors representing low co-expression and dark values representing high co-expression.

### Translatome analysis of spinal neurons in neuropathic mice

The modality specificity of sensory processing is severely compromised in chronic pain states, especially in those resulting from peripheral nerve damage [34–36]. We therefore decided to apply our bacTRAP approach to mice with neuropathic nerve lesions and to test whether peripheral nerve damage affects the neuron type-specific translatome. In the course of our above described experiments, we noticed that a lower mRNA yield (e.g. from VGAT::bacTRAP mice) correlated with a higher inter-sample variability in the number of detected reads per gene and in generally lower significance levels. In the initial experiments, we isolated the entire lumbar spinal cord (approx. L2-L6). In this second set of experiments we restricted our analysis to the lumbar segments L3 – L6 (i.e., to the segments innervated by the damaged sciatic nerve) of the ipsilateral spinal cord. Because of the smaller size of the tissue sample, we expected to have a more than 2 – 3-fold reduction in RNA yield after translating ribosome purification. To compensate for this smaller amount, we pooled three spinal cords per sample. To assess the reproducibility of the modified bacTRAP approach, we first compared the translatomes isolated from pooled spinal cords of naive VGLUT2::bacTRAP and Gad67::bacTRAP mice. We then compared the results to our initial analysis. We detected 13’033 genes of which 13’003 genes were also detected in our first analysis. Applying differential gene expression analysis, we detected 3180 significantly enriched genes (FDR £ 0.05) in this second compared to 1812 genes in the first analysis. Seventy percent of the genes (1269) enriched in the first analysis were also detected in the second analysis. Eighty-seven percent of the genes enriched in the first analysis and displaying a fold change ≥ 2 were also found to be enriched in the second analysis. Hence, we concluded that the approach is highly reproducible. The increase in the number of detected genes is likely due to a reduced inter-sample variability in the pooled samples.

We then focused on the analysis of VGLUT2::bacTRAP and Gad67::bacTRAP mice after employing two different types of nerve lesions (chronic constriction injury (CCI) [37] and spared nerve injury (SNI) of the sciatic nerve [38]. We characterized the translatome of the respective neuronal populations in mice that either underwent sham (surgery without constricting suture or lesion), CCI, or SNI surgery 7 days post-surgery when the neuropathy-induced hypersensitivity was fully established. We verified that all mice included in this analysis had developed mechanical hypersensitivity in response to the nerve injury.

We then compared gene expression differences between the three experimental groups of mice for glutamatergic and GABAergic neurons (first order comparisons) (Fig 7 and suppl. Information/Table S17). We only considered genes with normalized counts of ≥ 50 in at least two samples within any group. With sham-operated mice as the reference, DGEAs revealed significant changes in gene expression (FDR £ 0.05) caused by CCI and SNI surgery within both neuron populations (Fig 7). Four genes were up-regulated by SNI surgery in GABAergic (Gad67::bacTRAP) neurons relative to sham-operated mice. *Sprr1a* was the only gene up-regulated in GABAergic neurons after CCI surgery. Very similar results were obtained for the glutamatergic (VGLUT2::bacTRAP) neuron population. Three genes were up-regulated in SNI (versus sham) operated mice and one gene (*Sprr1a*) was up-regulated by CCI versus sham. No down-regulated genes were found in either the glutamatergic or the GABAergic population.

**Figure 7.**
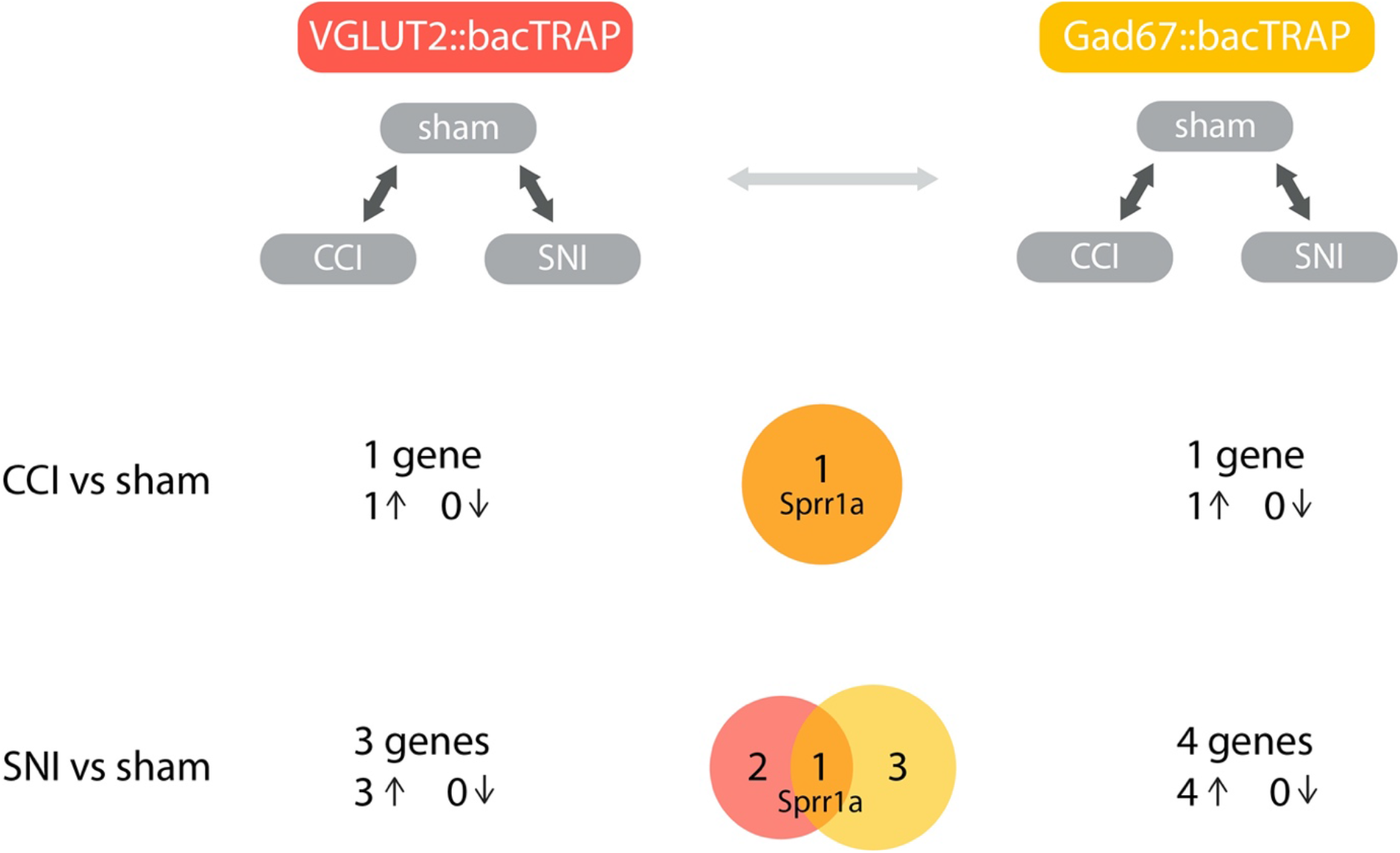
Differential gene expression in excitatory (VGLUT2::bacTRAP) and inhibitory (Gad67::bacTRAP) spinal cord neurons after peripheral nerve injury. Translatome analysis of Gad67::bacTRAP and VGLUT2::bacTRAP mice: differentially expressed genes in both mouse lines for the comparisons of CCI over sham and SNI over sham with number of significantly differentially expressed genes (FDR ≤ 0.05) and number of upregulated (arrow up ↟) and downregulated genes (arrow down ↓). Venn diagrams represent the intersection of differentially expressed genes for each comparison between the two mouse lines with number of genes exclusive to each group and common in between the two groups.

We found that *Sprr1a* expression in glutamatergic neurons displayed an approximately 25-fold upregulation after CCI and a 200-fold up-regulation after SNI (Fig 8A). *Sprr1a* was also up-regulated in Gad67-positive neurons after CCI and SNI (approximately 45-fold and 150-fold, respectively). Immunohistochemistry confirmed a strong up-regulation of Sprr1a protein in the ipsilateral dorsal horn and in ipsilateral motoneurons (Fig 8B). Staining in the dorsal horn was confined to the neuropil with no clear signal from neuronal somata, while in the ventral horn motor neuron somata were clearly labelled. Subsequent *in situ* hybridization experiments confirmed upregulated expression in motor neurons but failed to detect *Sprr1a* in neurons of the dorsal spinal cord (i.e., RNAscope signal around nuclei of spinal dorsal horn cells) (Fig 8F-J), suggesting that the signal in the dorsal horn detected in the immunohistochemistry experiments originated from DRG neuron axons and terminals. In conclusion, we find surprisingly little changes in the translatome of excitatory or inhibitory spinal neurons after peripheral nerve injury, suggesting that neuropathy-induced changes in spinal circuits may rely on alterations occurring independent of changes in neuronal translatomes.

**Figure 8.**
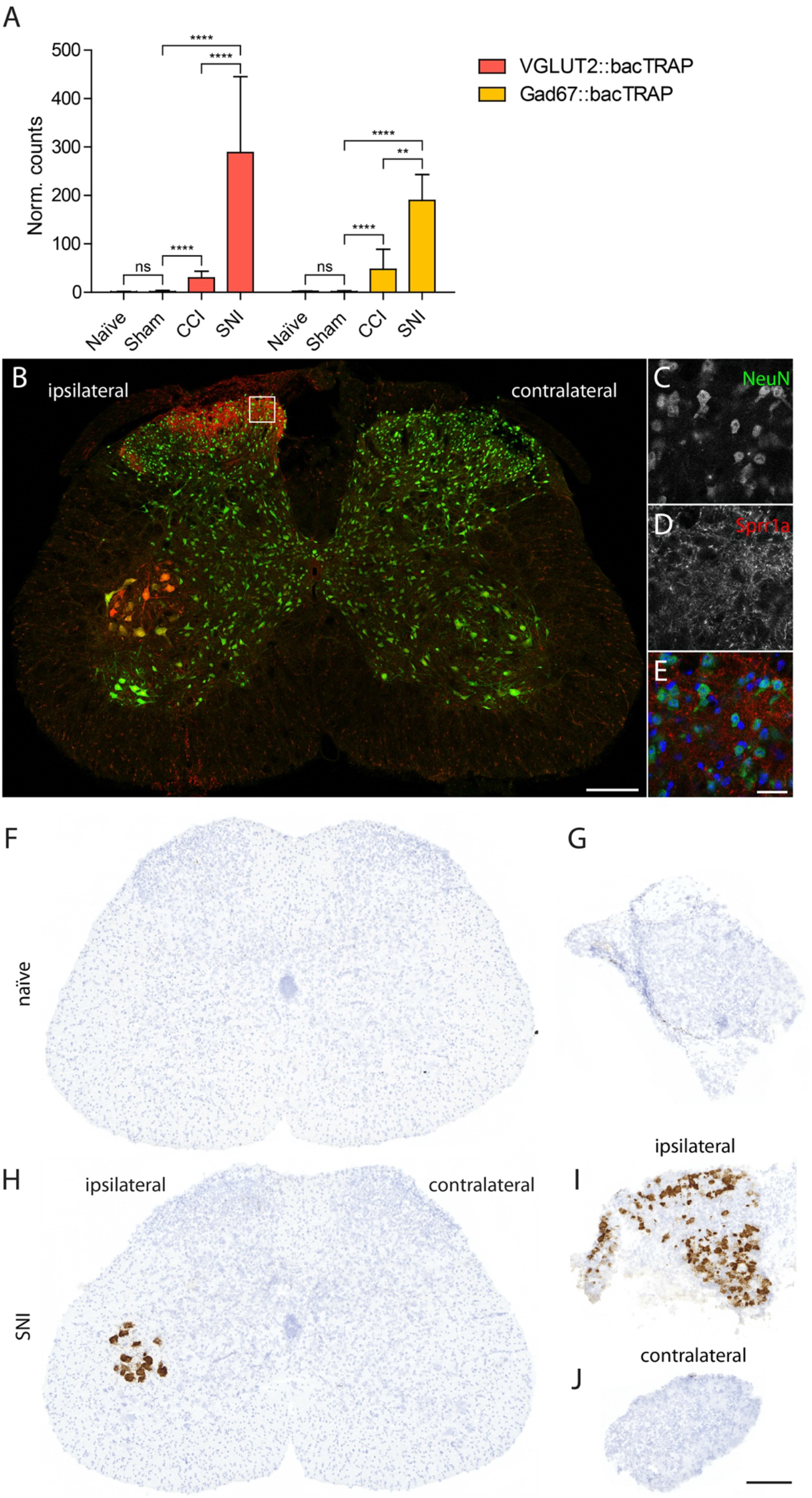
Upregulation of Sprr1a expression after peripheral nerve injury. (A) Translatome data of Sprr1a in naïve, sham, CCI and SNI operated VGLUT2::bacTRAP and Gad67::bacTRAP mice. Expression levels are significantly (p ≤ 0.05) increased in CCI and SNI as compared to sham surgery and in SNI compared to CCI in both mouse lines. (B-E) IHC against Sprr1a (red) and NeuN (green) in the transverse, lumbar spinal cord sections of a C57BL/6J mouse that underwent SNI surgery (seven days post-surgery). (B) Overview over the whole spinal cord sections, Sprr1a in red is mostly seen in fibers in the medial and lateral, superficial dorsal horn and in the soma of motoneurons in lamina IX on the ipsilateral side. Box indicated position of higher magnification image in (C-E). (C) NeuN, (D) Sprr1a and composite with nuclear DAPI staining in blue of medial, superficial dorsal horn with fibrous staining of Sprr1a. (F-J) ISH for Sprr1a in spinal cord (F, H) and DRGs (G, I, J) of naïve C57BL/6J mice (F-G) and C57BL/6J mice that underwent SNI surgery, seven days post-surgery (H-J). (I) DRG ipsilateral and (J) contralateral to the injured nerve. * p ≤ 0.05, ** p ≤ 0.01, *** p ≤ 0.001, **** p ≤ 0.0001, error: ± SD. Scale bar (B, F-J) 200 μm, (C-E) 20 μm.

## Discussion

Several groups have recently employed single cell or single nucleus sequencing to obtain a transcriptional atlas of the spinal cord [8, 9, 39]. We have used a complementary approach to specifically identify the mRNA translation profiles (translatomes) of inhibitory (GABA/glycinergic) and excitatory (glutamatergic) spinal cord neurons in naïve and neuropathic mice.

### Single cell transcriptome profiling versus cell type specific translatome profiling

mRNA translation profiles obtained by the TRAP approach provide cell-type-specific gene expression but not the single cell resolution obtained with single cell transcriptome profiling. On the other hand, tissue-dissociation procedures needed for single cell analyses often lead to undesired responses in gene expression and RNA degradation that are avoided when the bacTRAP technology is used. Furthermore, the fraction of polysome-bound mRNA should more accurately reflect protein levels than approaches based on nuclear or total RNA analysis [16]. The TRAP approach therefore represents a complimentary approach to bulk tissue and single cell sequencing capable to reveal novel insights into tissue diversity and pathological changes. In fact, two recent studies have employed the TRAP approach to primary nociceptors and uncovered unknown differences between trigeminal and dorsal root ganglia as well as unidentified driver genes for neuropathic pain [40, 41].

In our translatome comparison of inhibitory and excitatory spinal neurons, we find a high degree of overlap between genes that have been suggested to label subpopulations of neurons by single cell RNAseq experiments and those genes enriched in our analyses. However, we also find genes enriched in inhibitory and excitatory neurons that were either not detected in single cell RNAseq or where detection levels were too low to assign the gene to a subpopulation (e.g. *Phox2a*) or were potentially filtered out to reduce artefacts (e.g. cell stress responses) that might have been introduced by dissociation of the tissue to obtain single cells (e.g. *Ighg3*). Hence, the sensitivity of the TRAP approach appears at least equal to single cell RNA sequencing. A potential disadvantage of the TRAP approach is the spill-over of unbound RNA molecules from one neuron population to RNA free eGFP-tagged ribosomes of another neuron population. In fact, we detected low levels of most of the genes that are thought to be expressed exclusively in either inhibitory or excitatory neurons also in the respective other subset. Such examples include *Slc17a6* (VGLUT2), *Slc32a1* (VIAAT), *GAD1/2* or *Slc6a5* (Glyt2). These genes were detected at read numbers that correspond to 1 - 6% of the read numbers in the transgene expressing population. It is likely that the detection of *Sprr1a* in GABAergic and glutamatergic spinal neurons after either CCI or SNI resulted from spill-over from spinal motor neurons, in which *Sprr1a* is massively upregulated after peripheral nerve damage

### Translatome differences between dorsal horn neurons of naïve mice

Our GO term and pathway analyses of enriched translated genes revealed decisive differences between different subsets of spinal neurons in three gene families. We identified several transcription factors that are required during development for decisions about an excitatory and inhibitory cell fate but remain expressed into adulthood in a cell-type-specific manner. Although most neurotransmitters act both on excitatory and inhibitory neurons, we identified subtypes of neurotransmitter receptors and of down-stream signalling cascades that differ between excitatory or inhibitory neurons. Finally, we found that the heterogeneity within the large population of excitatory dorsal horn neurons is impressively mirrored by the expression of 11 neuropeptides or neuropeptide precursors.

### Transcription factors

Thirty to thirty-eight percent of the 50 genes most strongly enriched in excitatory or inhibitory neurons were transcription factors. Many of them have been described as crucial regulators during spinal cord development to establish cell type identity and thereby set up neuronal diversity [15]. Among these are genes such as *Olig3* which together with *Lbx1* (expressed in inhibitory and excitatory adult spinal neurons) are required to set up class A and class B neurons which populate the deeper or the upper laminae of the dorsal horn, respectively. Another battery of enriched genes such as those coding for the paired box factors (*Pax2/5/8*) and the Lim domain homeobox genes *Lhx1/5* on the one hand and *Tlx1/3 Lmx1b, Prrxl1, Phox2a* and others on the other hand are required to establish neurotransmitter identity and govern subsequent subtype specification [15]. While some of these genes have previously been reported to maintain expression into adult stages [8, 9, 22] many of them are believed to be switched-off in the adult [8, 15]. Our data however indicates sustained expression of the majority of these genes, suggesting also a hitherto unknown function in the adult.

### Differential use of neurotransmitter signaling pathways

Spinal inhibitory and excitatory neurons receive synaptic inputs from a wide variety of peripheral, local or supraspinal neurons, including glutamatergic, GABAergic, glycinergic, serotonergic, cholinergic, and adrenergic neurons. We find that both inhibitory and excitatory spinal neurons express receptors for all these neurotransmitters (Table S2–12). However, in some cases, neurotransmitter signaling appears to occur via different receptor subtypes and down-stream signaling cascades. Expression of several glutamate receptors of the kainate family (Grik1-3) is enriched in inhibitory spinal neurons. Kainate receptors display different modes of signaling and a metabotropic action of kainate receptors has been involved in presynaptic regulation and inhibition of GABA release [42, 43]. A preferential expression of kainate receptors on inhibitory spinal neurons might therefore be involved in a glutamate-mediated inhibition of GABA (and glycine?) release. Even more striking is the enrichment of several serotonin receptors on inhibitory neurons with the most pronounced example being Htr3a, the only ionotropic serotonin receptor. Expression of Htr3a is more than 8-fold enriched in inhibitory neurons and *in situ* hybridization indicated expression restricted to the deep dorsal horn. It is therefore conceivable that serotonin release from descending supraspinal neurons has a differential effect onto excitatory and inhibitory spinal neurons.

### Lamina specific expression of neuropeptides suggests a neuropeptide code involved in sensory perception

Our data confirm and extend previous single cell studies that identified neuropeptides as markers of excitatory neuronal subtypes [8, 9]. Our multiplex in situ hybridization experiments demonstrate a lamina-specific distribution of many of the enriched neuropeptides with rather limited expression overlap between different neuropeptides. In total we find that the 11 different neuropeptides analysed in this study span dorsal laminae I-V, thus setting up a map for a potential neuropeptide sensory signalling code. Indeed, emerging evidence suggests that the release of specific neuropeptides by distinct dorsal horn neurons is crucial for sensory perception. Signalling by gastrin releasing peptide (GRP), which is found concentrated in lamina II is critical for itch perception in mice [44–47] and required for successful signal transmission from second order GRP to third order GRP receptor neurons in lamina II of the spinal cord [48]. Another example is somatostatin (SST), which is also expressed in lamina II dorsal horn neurons. After release from peripheral sensory neurons or spinal dorsal horn neurons it inhibits inhibitory dynorphin (Dyn) neurons, which in turn inhibit GRP receptor expressing neurons [49]. SST release thereby facilitates pruritoceptive signalling. In this context, it is interesting to note that the majority of spinal SST expressing neurons are excitatory (glutamatergic), while the effect of SST on Dyn neurons is inhibitory. These studies demonstrate that neuropeptides are able to add an additional layer of complexity to neuronal communication and may in certain cases dominate over the effect of fast-acting neurotransmitters such as glutamate, GABA and glycine. The lamina specific expression of many neuropeptides in dorsal horn neurons suggests that distinct groups of neuropeptide expressing cells are activated by a restricted subset of primary afferents. Hence, distinct neuropeptides might only be released upon specific sensory stimuli. Our expression matrix provides a basis for further studies to decipher the function of neuropeptide expressing cells and of neuropeptide release in sensory signalling.

### No overt changes in the translatome of spinal neurons after peripheral nerve injury

It is generally accepted that altered somatosensory and nociceptive perception in neuropathic pain result from still incompletely understood changes at the dorsal horn circuitry [50–52]. Several of the suspected changes occur in second order intrinsic dorsal horn neurons. Our analyses in two types of peripheral nerve lesions failed to reveal major alterations in the neuronal translatome. The only gene consistently up-regulated in both neuropathic pain models was *Sprr1a*. Interestingly, *Sprr1a* appeared up-regulated mainly, if not exclusively, in motor neurons and primary sensory DRG neurons, indicating that it became expressed in the injured neurons rather than in intrinsic order dorsal horn neurons. This up-regulation is consistent with its well-established role in axon repair [53, 54].

At a first glance, the low number of regulated genes detected in our study appears to be in stark contrast with previous work. We see at least two possible explanations. First, these previous studies analysed total RNA from the entire spinal cord [55], i.e., not only neuronal RNA but also RNA from astrocytes and microglia. These non-neuronal cells are major contributors in the central sensitization process following peripheral nerve damage [56–58]. Indeed, previous studies analysing whole spinal cord RNA detected the most profound and consistent changes in genes related to immune cell function [55, 58]. Second, a key difference between the present and previous studies is the use of the TRAP approach in our study, which limits the analysis to transcripts that are bound by the ribosome and will therefore likely be translated into proteins. Our data thus may suggest that many transcriptional changes are not propagated to the translatome. Several previous studies have indeed provided evidence that transcriptional changes do not necessarily translate into changes of the proteome [59–62]. Alternatively, some neuropathy-induced changes in the translatome might have been missed because they were transient. However, previous transcriptomics studies observed the most pronounced changes in gene expression at 7 days post-surgery, when the phenotype was fully established [55]. Finally, we cannot completely exclude that the critical translatome changes occurred in neuron population too small to be detectable in our samples of excitatory and inhibitory neurons. Despite these potential limitations, our data still suggest that neuropathy-induced circuit changes occur independent of major changes of the translatome of dorsal horn neurons.

## Materials and Methods

### Generation of VGLUT2::bacTRAP, VGAT::bacTRAP and Gad67::bacTRAP mouse lines

The VGLUT2::bacTRAP mouse line was generated by placing cDNA for the eGFP-L10a (TRAP) transgene, followed by the bovine growth hormone polyadenylation (poly(A)) signal in frame into exon 2 of the VGLUT2 (*Slc17a6*) gene (on BAC clone RPE23-84M15), using homologous recombination in bacteria [63], replacing most of exon 2 of the VGLUT2 gene. The start codon (ATG) in exon 1 was mutated to an XhoI restriction site. For the VGAT::bacTRAP mouse line the transgene was placed in frame into the end of exon 1 of the VGAT (*Slc32a1*) gene, replacing it partially (on BAC clone RPE23-392P11). For the Gad67::bacTRAP mouse line the transgene was placed in frame into the end of exon 2 of the Gad67 (*Gad1*) gene, replacing it partially (on BAC clone RP24-256C12). BAC DNA was injected into C57BL/6 oocytes. The thereby generated mice were screened for the presence of the eGFP-L10a transgene and for the two BAC ends by PCR. The transgenic mouse lines were maintained on a C57BL/6J (Black 6, The Jackson Laboratory, Stock No: 000664) background. C57BL/6J mice were also used as wild type controls. Mouse breeding and maintenance were provided by the Laboratory Animal Services Center (LASC) of the University of Zurich. Mice received food and water ad libitum and were kept in a 12 h light/dark cycle.

### Genotyping PCRs

The presence of the eGFP-L10a transgene in the three different mouse lines was determined using a specific PCR for the transgene or a generic PCR against eGFP with one of the following primer pairs:

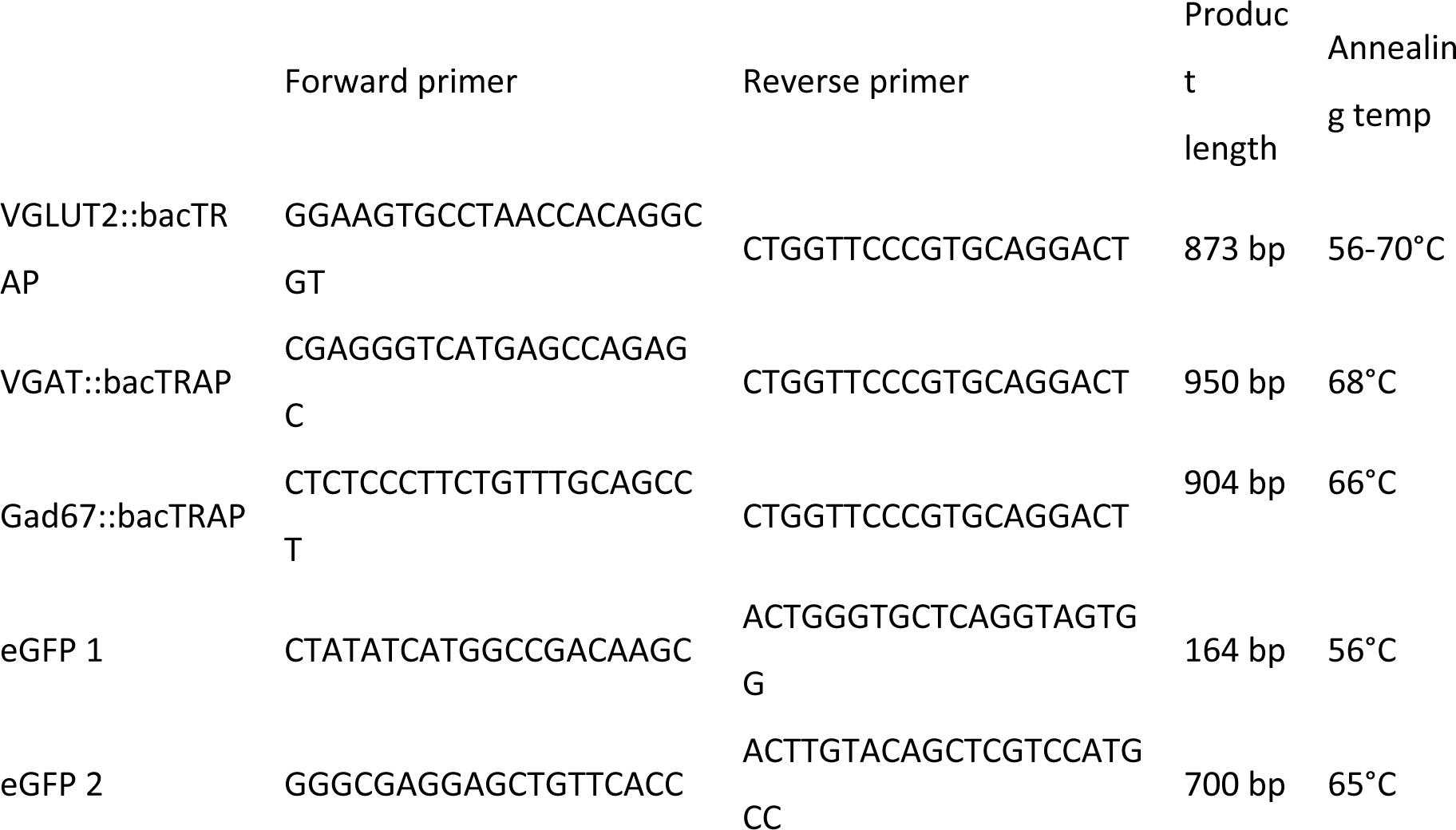

### Immunohistochemistry (IHC)

For mouse line validation, adult (> 2 months old), male and female bacTRAP mice and for the Sprr1a staining, adult (10-12 weeks old) male C57BL/6J mice, seven days after SNI surgery were used. Mice were transcardially perfused with approximately 20 ml of ice-cold artificial cerebrospinal fluid (ACSF, pH 7.4) or phosphate buffered saline (PBS, pH 7.4), followed by 100 mL of 2 or 4 % ice-cold paraformaldehyde (PFA, in PBS or 0.1 M Sodium phosphate buffer (PB), pH 7.4). The lumbar spinal cord was dissected and post-fixed for 1.5 - 2 h in 4 % PFA solution (in PBS or 0.1 M PB, pH 7.4), followed by incubation in 30 % sucrose (in PBS or 0.1 M PB, pH 7.4) for cryoprotection at 4°C overnight. Cryoprotected spinal cords were embedded in NEG50 frozen section medium (Richard-Allen Scientific) and stored at - 80°C until cutting into 25 μm thick sections on a Hyrax C60 Cryostat (Carl Zeiss). Sections were mounted on Superfrost Plus glass slides (Thermo Scientific, Zurich, Switzerland) and stored at −80°C.

For immunofluorescence staining the slides were washed for 5 min in PBS to remove the embedding medium, followed by blocking with 5 % normal donkey serum (NDS, AbD Serotec, RRID:SCR_008898) in 0.1 % Triton X-100-PBS for at least 30 min at room temperature (RT). Sections were incubated with primary antibodies (see resource table) in the blocking solution at 4°C overnight, followed by three washes in PBS for 5 min and incubation with secondary antibodies (fluorophore-coupled donkey antibodies, Jackson ImmunoResearch) in blocking solution at RT for 30 to 60 min. Subsequently, sections were washed in PBS and incubated with 4’,6-diamidino-2-phenylindole (DAPI, 500 ng/ml in PBS) for 10 min and again washed in PBS. Finally, sections were covered with DAKO fluorescent mounting medium (Dako, RRID:SCR_013530) and coverslips.

### *In situ* hybridization (ISH)

For the Sprr1a-ISH, 6.5 weeks old, male C57BL/6J mice were sacrificed seven days after SNI surgery. For all other ISHs, 6 weeks old, naïve male C57BL/6J mice were used. For spinal cord preparation, the vertebral column containing the lumbar part of the spinal cord were dissected, immediately after decapitation of the mouse. A syringe filled with ice-cold, RNAse-free PBS was fitted tightly against the caudal opening of the spinal canal and pressure was applied to press the spinal cord out, which was timed to the lumbar part. DRGs were prepared from the vertebral column containing the lumbar DRGs, which was cut open from the ventral side using fine scissors. After dissection, spinal cords were snap frozen in liquid nitrogen and stored at −80°C until embedding in NEG50 frozen section medium (Richard-Allen Scientific). DRGs were embedded directly after preparation. Frozen blocks were again stored at −80°C until cutting into 12 or 16-25 μm thick sections for DRGs and spinal cords, respectively and mounted as described for IHC.

For single-plex chromogenic ISH the manual RNAScope^®^ 2.5 HD BROWN Assay (ACD, Bio-Techne, catalog no. 322300) was used. The manufacturer’s pretreatment protocol for fresh frozen tissue (document no. 320536-TN, rev. date 11112016) and detection protocol (document no. 322310-USM, rev. date 11052015) were followed. Hybridization with Amp 5 was increased from 30 to 45 min for Sprr1a and to 60 min for all other probes in order to increase the signal. Counterstaining in 50 % hematoxylin was reduced from 2 min to 30 sec. Probes are listed in the resource table.

For multiplex fluorescent ISH (FISH) the manual RNAScope^®^ Multiplex Fluorescent Assay (ACD, Bio-Techne, catalog no. 320850) was used. The manufacturer’s pretreatment protocol for fresh frozen tissue (document no. 320513, rev. date 11052015) and detection protocol (document no. 320293-UM, rev. date 03142017) were followed. The fluorophore alternatives (Amp 4 Alt) was chosen in such a way, that when possible the weakest expressing gene would lie in red channel (Atto 550) and not in the far-red channel (Atto 647). The 3-plex negative control probe was amplified with the corresponding Amp 4 Alt. Probes are listed in the resource table. For every neuropeptide combination, one slide with four lumbar spinal cord sections of four animals, each, was reacted.

### Image acquisition and analysis

Bright field imaging of ISHs and fluorescent imaging of FISHs and IHCs was done using the following microscopes:

| Microscope | Objectives | Software | Purpose |
| --- | --- | --- | --- |
| Zeiss Axio Scan.Z1 slidescanner | 5x/0.25 Fluar air<br>10x/0.45 Plan-Apochromat air | ZEN 2 slidescan (blue edition) | Bright field images of chromogenic ISHs |
| Zeiss LSM710 confocal microscope | 20x/0.8 Plan-Apochromat air<br>40x/1.3 EC Plan-Neofluar oil | ZEN 2011 (black edition) | Fluorescent images for mouse line validation and Sprr1a IHC staining |
| Zeiss LSM700 confocal microscope | 25x/0.8 Plan-Neofluar oil<br>40x/1.4 LCI Plan-Apochromat oil | Zen 2011 (black edition) | Overview images of multiplex FISH |
| Zeiss LSM 800 with Airyscan / confocal | 25x/0.8 Plan-Neofluar oil<br>40x/1.4 Plan-Apochromat oil | ZEN 2.3 (blue edition) | Overview and analysis images of multiplex FISH |

For fluorescent imaging, the pinhole was set to 1 airy unit for every channel, which were scanned sequentially to avoid overlapping emission spectra or with a combination of the ultraviolet and infrared channel in one track, where emission spectra overlap is minimal.

For the validation of the correct expression of the eGFP-L10a transgene, the 40x objectives on the LSM 700 or 710 were used with a zoom of 0.6x to acquire a tile scan over the complete lumbar dorsal horn (Pixel size: x 0.346 μm, y 0.346 μm, z 1.628/2.487 μm). A small z-stack (usually, 3.3 μm within 3 optical sections) at a bit depth of 8 was acquired. For the analysis, the cell counter plugin of ImageJ was used. Manual counting was done in the middle optical section; the other sections were used as an aid to distinguish between neighboring cells. Three hemi-sections from three animals of each mouse line were analyzed. Ratios were calculated per animal and then averaged.

For the Sprr1a IHC staining, overview images of the dorsal horn were taken using tile scans with the 20x objective (Pixel size: x 0.415 μm, y 0.415 μm). Detail images were taken with the 40x objective (Pixel size: x 0.104 μm, y 0.104 μm, z 0.329 μm).

For the quantification of the co-expression of the neuropeptides with multiplex FISH, the 40x objective on the LSM 800 was used with a zoom of 2x and an image size of 1024 x 1024 px (pixel size: x 0.078 μm, y 0.078 μm). An array of non-overlapping images that covers the complete dorsal horn was scanned at a bit depth of 16. Analysis was done with CellProfiler 2.2.0 software [64]. The cell profiler pipeline can be provided upon request. Briefly, nuclei were detected using the *IdentifyPrimaryObjects* module and expanded with the *IdentifySecondaryObjects* module to define the cell soma (from now on referred to as “cell”). The signal dots were detected using the *IdentifyPrimaryObjects* module, adapting the “threshold correction factor” and the “lower and upper bounds on threshold” for every slide (neuropeptide combination) to minimize false positive and false negative detection. The signal dots of the different channels were assigned to the cells they lie in using the *RelateObjects* module. The number of cells and related signal dots (per channel) were exported using the *ExportToSpreadsheet* module. Images with artefacts or high background were manually excluded from the automatic analysis in order to avoid false positive signal. Per slide (neuropeptide combination), three animals with two hemi-sections per animal were analyzed. Additionally, per combination one hemi-section hybridized with the 3-plex negative control probe and amplified with the same Amp 4 Alt was imaged with the same microscope setting and analyzed with the same CellProfiler setting in order to determine the background.

Data analysis was conducted with R. To determine the signal background, the 3-plex negative control was analyzed. For every combination and channel, the 90^th^ percentile of dots per cell in the negative control was determined. This value was set as the threshold above which a cell is counted as positive for the respective signal. 20 dots per cell were set as a threshold for high expression. The number of single, double and triple-positive cells was counted. Ratios were calculated per animal and then averaged.

### Mouse models for neuropathic pain using peripheral nerve injury

Mice were anesthetized using isoflurane (5 min induction chamber with 3-5 % isoflurane, maintenance with 1-3 %). The isoflurane dose was adjusted to a level where the animal does not respond to pinching of its hind paw. Mice were placed on a heat mat to maintain body temperature. Lubricant eye ointment was applied to prevent corneal drying during surgery. 0.1-0.2 mg/kg buprenorphine were injected subcutaneously as analgesic treatment. The left hind limb was shaved with an electric razor and the skin was swabbed with iodine solution and allowed to dry. A longitudinal cut of 10-15 mm was made to the lateral surface of the thigh and a section was made through the biceps femoris muscle exposing the sciatic nerve and its three branches: the tibial, common peroneal and the sural nerve. For chronic constriction injury (CCI), three loose ligatures were placed around the nerve with 5-0 surgical silk proximal to its trifurcation [65]. For spared nerve injury (SNI), the common peroneal and the tibial nerve were transected distal to trifurcation of the nerve, removing 2 mm. The sural nerve was left intact [66]. For sham surgery the sciatic nerve was exposed but not injured. The surgical wound was closed in two layers (muscle and skin). 7 days postsurgery, when thermal and mechanical sensitization are fully established, animals were tested for mechanical hypersensitivity using von Frey filaments. Afterwards animals were sacrificed for immunopurification of polysomal mRNA or tissue preparation for IHC or ISH.

### Behavioral testing

Measurements were done before surgery (baseline) and 7 days post-surgery. Withdrawal thresholds to a static innocuous mechanical stimulus were assessed using an electronic **von Frey** anesthesiometer with a 7 g filament (IITC, Woodland Hills, CA). Mice were kept in a 10 x 10 cm Plexiglas box, sitting on an elevated grid platform. The lateral plantar surface of the hind paw (innervated by the sural nerve) was poked with the filament and the force at which the mouse retracted its paw was measured. Each paw was measured 7-8 times with intervals of at least 8-10 min.

### Cell-type-specific polysomal mRNA isolation from excitatory and inhibitory spinal cord neurons

To specifically isolate polysomal mRNA from genetically targeted cells, the three bacTRAP mouse lines described above were used and the protocol published by Heiman, Kulicke (67) was followed. In brief, solutions and the affinity matrix were prepared as described. For the affinity matrix of Gad67::bacTRAP 300 μl and of VGLUT2::bacTRAP and VGAT::bacTRAP 150 μl Streptavidin MyOne T1 Dynabeads and the corresponding volumes of biotinylated protein L and GFP antibodies 19C8 and 19F7 were used in a final volume of 200 μl.

In the comparison of excitatory to inhibitory neurons, three male mice per mouse line, aged between 8 and 10 weeks were used. For the analysis of changes after neuropathic pain, nine male mice per group, aged between 8 and 10 weeks were sacrificed 7 days post-surgery. The vertebral column containing the lumbar part of the spinal cord was dissected, immediately after decapitation. A syringe filled with ice-cold dissection buffer was fitted tightly against the caudal opening of the spinal canal and pressure was applied to press the spinal cord out, which was trimmed to the lumbar part. For the neuropathic pain analysis, the lumbar spinal cord was bisected along the midline and the left side (ipsilateral to the peripheral nerve surgery) was further processed. The tissue was washed, homogenized (3 halves pooled for the neuropathic pain analysis), centrifuged, lysed and again centrifuged as described in the protocol by Heiman et al. [67].

Immunopurification was done as described by Heiman et al. [67]. RNA purification was done using the RNeasy Plus Micro Kit (Qiagen, catalog no. 74034). RNA was eluted from beads with 350 μl Buffer RLT Plus from the kit, supplemented with 40 μM dithiothreitol (DTT, Thermo Fisher Scientific, catalog no. R0861) and then processed according to the manufacturer’s protocol. RNA concentration was determined using the RiboGreen fluorescence assay (Thermo Fisher Scientific, catalog no. R11491) on a NanoDrop 3300 microvolume fluorospectrometer. RNA quality was assessed by high sensitivity RNA screen tape on an Agilent 2200 TapeStation. RNA samples with RINe values of 6.8 to7.8 (inhibitory – excitatory comparison) and 6.0 to7.3 (neuropathic pain analysis) were used for sequencing library construction.

### Sequencing library construction, next-generation sequencing (NGS), mapping and differential gene expression analysis (DGEA)

The sequencing library was prepared using the Ovation Mouse RNA-Seq System 1-16 kit (NuGEN, catalog no. 0348-32). The manufacturer’s protocol was followed. 13-20 ng RNA were used as input material for the inhibitory – excitatory comparison and 30 ng RNA for the neuropathic pain analysis. The optional cDNA fragmentation step was performed. As recommended, the optimal cycle number for amplification was determined by qPCR.

Sequencing of the library, mapping to the reference genome, as well as DGEA were performed by the Functional Genomics Center Zurich (FGCZ). For the excitatory – inhibitory comparison, NGS was done on two lanes of a HiSeq 2500 system (illumina) with single-end sequencing of 125 bp. Mapping was done in R using the STAR package [68]. DGEA was done in R using either the edgeR or the DESeq2 package [69–71]. For the neuropathic pain analysis, NGS was done on two lanes of a HiSeq 4000 system (illumina) with paired-end sequencing of 2×150 bp. Mapping and DGEA were done in R using the packages STAR [68] and DESeq2, respectively [71]. In order to compare DGEAs of naïve mice (Gad67 vs VGLUT2) from the first analysis (entire spinal cord segments L2-L6) and the second analysis (spinal hemisegments from L3-L6) the DESeq2 package was used for the DGEAs.

### Gene Ontology (GO) term and pathway enrichment analyses

GO term enrichment analysis was done using the online PANTHER tool (http://pantherdb.org/). Genes significantly (FDR ≤ 0.05) and ≥ 2-fold enriched in excitatory (VGLUT2) over inhibitory (VGAT or Gad67) and vice versa were analyzed. In the PANTHER overrepresentation test, the sets of enriched genes were compared to the set of all expressed genes as a reference. GO molecular function (MF) complete was used as the annotation data set. The significantly overrepresented (FDR ≤ 0.05) MFs were analyzed. When several GO terms of a hierarchy were significantly overrepresented, only the most specific GO term was displayed in the graphical representation. For the analysis of Gad67 vs. VGLUT2, two redundant GO terms were not displayed in the graphical representation in order to allow comprehensibility.

Pathway analysis was done with Ingenuity Pathway Analysis (IPA, QIAGEN). Genes significantly (FDR ≤ 0.05) and ≥ 2-fold enriched in excitatory (VGLUT2) over inhibitory (VGAT or Gad67) and vice versa were analyzed. To get a coarse classification of the enriched genes into subcellular location and function (“type”), these parameters were exported from the software.

### Resources

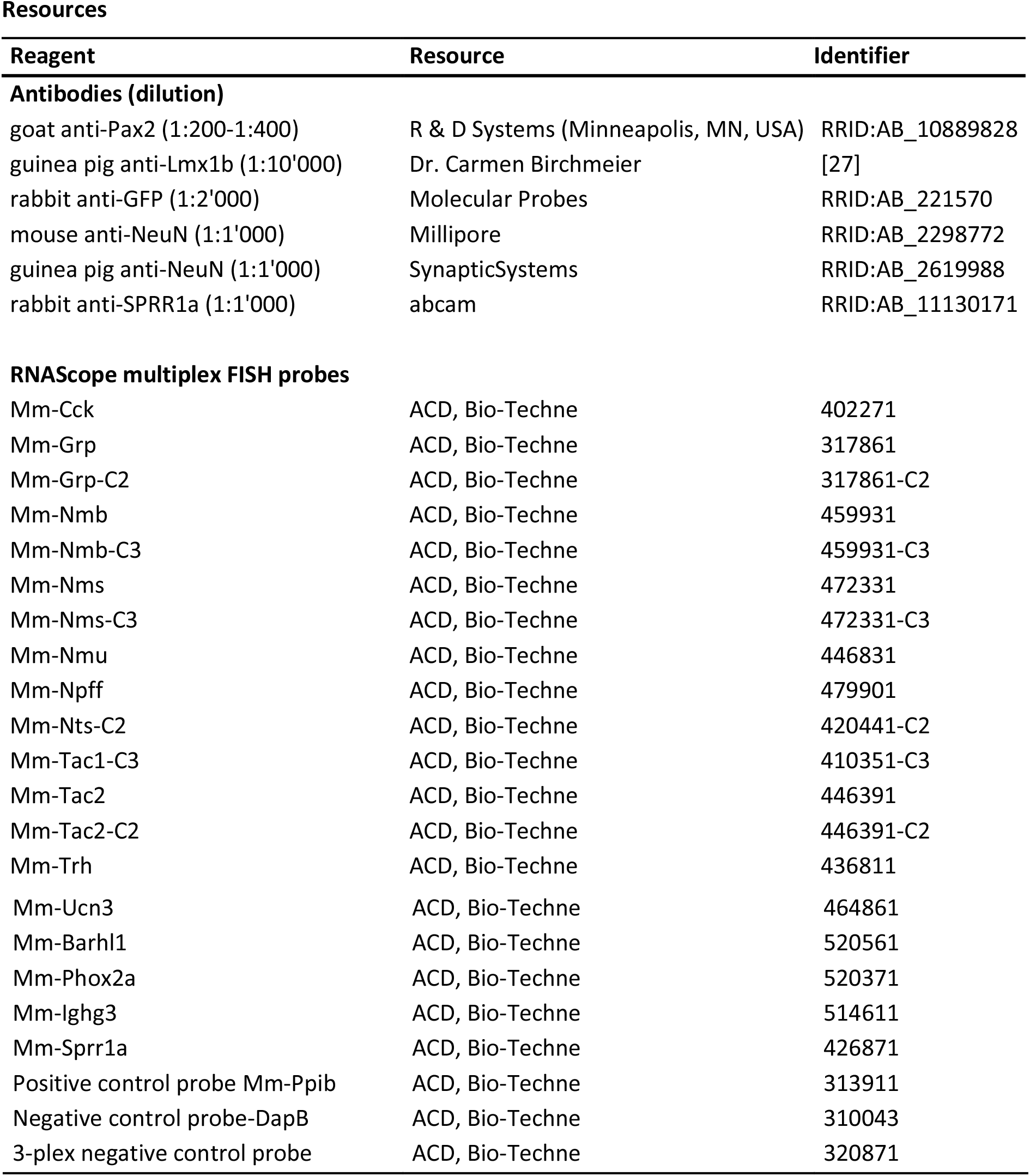

## Acknowledgement

The Functional genomics center Zurich (FGCZ) conducted next-generation sequencing, mapping, as well as DGEA. The FGCZ also assisted with data analysis. We are grateful to C. Birchmeier for providing the Lmx1b antibody. We thank Isabelle Kellenberger for genotyping the mice. This work was supported by grants from the Swiss National Science Foundation (SNSF, grant number 176398), an Advanced Investigator Grant from the European Research Council (ERC, grant number AdvG 250128) and a grant from the Swiss Federal Institute of Technology (ETH Fellowship) to H.U.Z and by a SNSF grant (grant number: CRSK-3_190622) to H.W.

---

**Figure S1.**
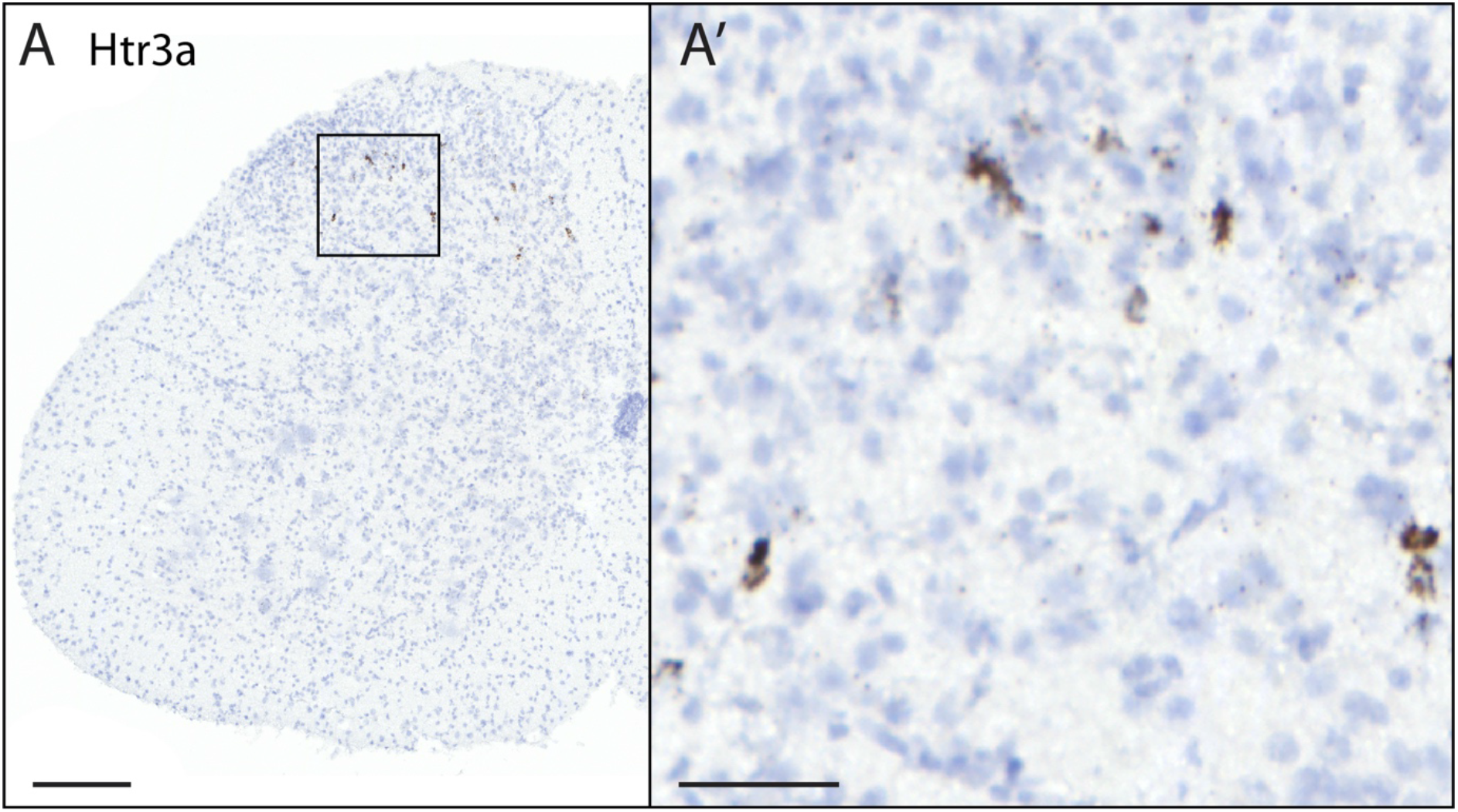
Expression pattern analysis of the ionotropic serotonin receptor Htr3a. Expression pattern analysis by *in situ* hybridization using a RNAscope probe directed against the mRNA encoding *Htr3a.* Scale bar: (A) 200 μm, (A’) 50 μm

**Figure S2.**
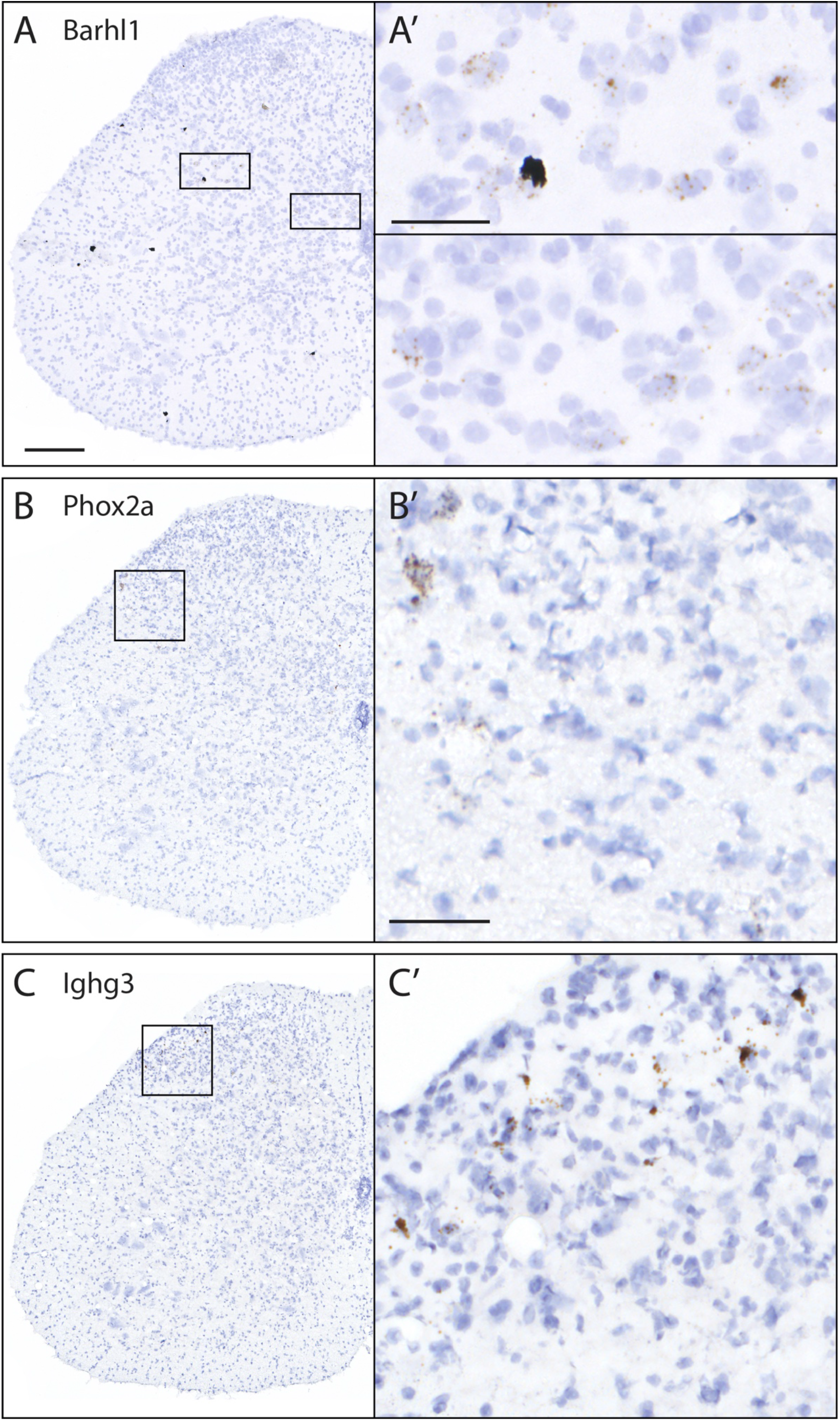
Confirmation of expression of genes with low detection levels, that were not detected in single-cell sequencing by Häring et al. [8]. *In situ* hybridization using RNAscope probes directed against (A) transcription factor BarH-like 1 (*Barhl1*), (B) transcription factor paired-like homeobox 2a (*Phox2a*), (C) Immunoglobulin heavy constant gamma 3 (*Ighg3*). Scale bar: (A, B, C) 200 μm, (A’, B’, C’) 50μm

## Most enriched genes

**Table S2.** Most enriched genes in vGluT2 vs. vGAT. Group: TF = transcription factor, NP = neuropeptide, pNP = putative neuropeptide (classification of neuropeptides and putative neuropeptides is based on Burbach (2011)), NF = neurotrophic factor, NPR = neuropeptide receptor. sc data = single-cell data from Haring et al. (2018): nd = not detected, ng = not grouped.

|  |  | vGluT2<br>[norm. count] | vGAT<br>[norm. count] | Gad67<br>[norm. count] | vGluT2 vs. vGAT |  | vGluT2 vs. Gad67 |  |  |
| --- | --- | --- | --- | --- | --- | --- | --- | --- | --- |
| Gene | Group | Mean ± Stddev | Mean ± Stddev | Mean ± Stddev | log2 Ratio | FDR | log2 Ratio | FDR | sc Data |
| Ucn3 | NP | 106 ± 34 | 0 ± 0 | 5.9 ± 10.2 | 9.87 | 8.52E-14 | 4.18 | 4.60E-04 | |
| Lhx2 | TF | 199 ± 35 | 0.96 ± 1.67 | 0 ± 0 | 7.62 | 9.29E-26 | 10.71 | 2.02E-35 | ng |
| Olig3 | TF | 176 ± 28 | 1.56 ± 1.86 | 12.2 ± 7.1 | 6.58 | 5.80E-20 | 3.82 | 4.92E-13 | nd |
| Barhl1 | TF | 382 ± 38 | 4.3 ± 2.9 | 0 ± 0 | 6.40 | 3.25E-49 | 11.65 | 5.24E-74 | nd |
| Shox2 | TF | 765 ± 32 | 11.5 ± 10.4 | 25.6 ± 20.9 | 6.12 | 7.88E-35 | 4.91 | 9.81E-21 | nd |
| Crh | NP | 219 ± 44 | 3 ± 2.6 | 0 ± 0 | 6.05 | 5.17E-24 | 10.84 | 4.50E-39 | |
| Barhl2 | TF | 480 ± 92 | 6.7 ± 7.8 | 15.4 ± 2.38 | 6.02 | 9.86E-37 | 4.94 | 7.52E-53 | nd |
| Slc17a6 | | 47533 ± 1973 | 753 ± 360 | 748 ± 191 | 5.98 | 6.32E-51 | 5.98 | 2.37E-110 | |
| Tlx3 | TF | 736 ± 64 | 13.7 ± 19.3 | 10.6 ± 9.6 | 5.78 | 1.63E-15 | 6.07 | 7.54E-69 | |
| Mgp | | 216.3 ± 16.7 | 4.5 ± 7.8 | 15 ± 16.2 | 5.60 | 4.43E-12 | 3.82 | 1.87E-08 | ng |
| Adcyap1 | NP | 2469 ± 241 | 52 ± 32 | 73.3 ± 17.2 | 5.59 | 2.04E-42 | 5.07 | 8.12E-134 | |
| Nrn1 | NF | 20203 ± 1915 | 457 ± 164 | 458 ± 49 | 5.47 | 8.16E-86 | 5.45 | 1.18E-207 | |
| Npff | NP | 317 ± 26 | 7.7 ± 10.9 | 7.9 ± 7.1 | 5.43 | 6.03E-15 | 5.34 | 1.38E-33 | |
| Pou4f1 | TF | 1462 ± 131 | 33.9 ± 21.8 | 43 ± 10.3 | 5.40 | 7.59E-50 | 5.09 | 1.94E-113 | |
| Ctla2a | | 104 ± 25 | 2.2 ± 3.8 | 8.1 ± 7.1 | 5.37 | 7.48E-07 | 3.69 | 3.89E-05 | ng |
| Lhx9 | TF | 214.8 ± 23.5 | 5.1 ± 6.5 | 1.29 ± 1.56 | 5.37 | 6.79E-17 | 7.23 | 2.13E-34 | nd |
| Pla2g5 | | 507 ± 73 | 12.7 ± 22 | 16.2 ± 6.7 | 5.33 | 1.84E-05 | 4.98 | 9.34E-52 | |
| Cacna2d1 | | 6755 ± 656 | 170.2 ± 82 | 203 ± 46 | 5.32 | 8.04E-77 | 5.05 | 4.24E-175 | |
| Vsx2 | TF | 1501 ± 91 | 41.9 ± 27.1 | 38.1 ± 6 | 5.17 | 6.07E-53 | 5.29 | 1.09E-132 | nd |
| Nms | NP | 217 ± 47 | 6.4 ± 11 | 1.48 ± 1.76 | 5.13 | 2.90E-07 | 6.96 | 2.35E-32 | |
| Tmem163 | | 4153 ± 563 | 122 ± 69 | 118 ± 13.3 | 5.10 | 1.45E-45 | 5.12 | 3.92E-176 | ng |
| Phox2a | TF | 136.9 ± 12.6 | 3.9 ± 6.7 | 3.4 ± 5.9 | 4.99 | 6.60E-07 | 5.17 | 4.54E-10 | nd |
| Bdnf | NF | 1097 ± 70 | 44 ± 36 | 18.5 ± 11.9 | 4.63 | 4.02E-15 | 5.88 | 9.28E-90 | |
| Nmur2 | NPR | 588 ± 84 | 24.9 ± 22.6 | 46 ± 38 | 4.59 | 6.13E-21 | 3.67 | 5.66E-16 | |
| Cbln1 | pNP | 6341 ± 1471 | 310 ± 95 | 322 ± 53 | 4.36 | 1.17E-69 | 4.29 | 4.46E-110 | |
| Nmu | NP | 571.9 ± 23.9 | 29 ± 34 | 7.6 ± 5.1 | 4.31 | 5.78E-06 | 6.17 | 3.25E-73 | |
| Chrm5 | | 177 ± 28 | 8.8 ± 8.4 | 23.1 ± 13.1 | 4.27 | 3.14E-09 | 2.91 | 7.15E-09 | |
| Nmb | NP | 204.9 ± 15.7 | 10.5 ± 8.2 | 4.5 ± 4.1 | 4.23 | 1.87E-12 | 5.51 | 1.08E-23 | |
| Tac2 | NP | 5416 ± 363 | 299 ± 86 | 536 ± 36 | 4.18 | 2.90E-85 | 3.33 | 7.35E-131 | |
| Nts | NP | 2582 ± 331 | 143 ± 86 | 130.6 ± 5.9 | 4.18 | 9.72E-31 | 4.29 | 9.74E-125 | |
| Lmx1b | TF | 1745 ± 90 | 99.1 ± 24.9 | 62.9 ± 9.1 | 4.14 | 9.36E-78 | 4.77 | 2.02E-129 | nd |
| Grpr | NPR | 292 ± 34 | 17.2 ± 22 | 26.7 ± 11.6 | 4.04 | 6.37E-07 | 3.42 | 3.11E-21 | |
| C1ql3 | | 2262 ± 56 | 140 ± 22 | 171 ± 57 | 4.00 | 3.75E-94 | 3.71 | 5.63E-61 | |
| Cpne4 | | 2929 ± 473 | 186.8 ± 7.8 | 300 ± 40 | 3.97 | 1.07E-89 | 3.28 | 8.89E-71 | |
| Lhx4 | TF | 249.1 ± 21.4 | 16.8 ± 17.9 | 25.3 ± 9.5 | 3.90 | 3.62E-06 | 3.29 | 2.32E-18 | nd |
| Slc17a8 | | 186 ± 28 | 12.5 ± 9.3 | 35.4 ± 15.6 | 3.86 | 2.31E-09 | 2.39 | 1.94E-07 | ng |
| Cck | NP | 7261 ± 778 | 538 ± 152 | 373 ± 74 | 3.76 | 4.13E-68 | 4.27 | 2.02E-140 | |
| Mafa | TF | 603 ± 78 | 46 ± 26 | 54.3 ± 40.2 | 3.71 | 3.26E-20 | 3.46 | 2.32E-17 | nd |
| Prrxl1 | TF | 1436 ± 65 | 124 ± 59 | 180.8 ± 23.9 | 3.54 | 5.86E-34 | 2.98 | 9.05E-61 | |
| F2rl2 | | 2146 ± 380 | 187 ± 156 | 264.1 ± 37.2 | 3.52 | 1.68E-12 | 3.01 | 1.04E-56 | |
| Ebf2 | TF | 3619 ± 436 | 355 ± 56 | 894.5 ± 59.7 | 3.35 | 8.47E-71 | 2.01 | 1.66E-40 | |
| Aldh1a2 | | 179 ± 75 | 17.6 ± 11.8 | 17 ± 14.2 | 3.35 | 1.42E-07 | 3.35 | 1.94E-07 | |
| Serpina3g | | 148 ± 36 | 15.8 ± 16.7 | 40.2 ± 8.1 | 3.23 | 8.13E-05 | 1.87 | 1.01E-04 | nd |
| Nptx2 | | 1325 ± 141 | 146.7 ± 24.5 | 181.2 ± 5.5 | 3.17 | 5.94E-48 | 2.86 | 1.79E-54 | |
| Grp | NP | 1085 ± 211 | 127 ± 39 | 105 ± 35 | 3.10 | 1.25E-28 | 3.36 | 2.48E-36 | |
| Car12 | | 423.4 ± 10.1 | 52.2 ± 9.1 | 71 ± 70 | 3.02 | 1.25E-23 | 2.56 | 1.64E-04 | |
| Pou4f2 | | 207 ± 39 | 25.7 ± 13.6 | 8.5 ± 4.4 | 3.01 | 1.79E-09 | 4.61 | 8.21E-21 | ng |
| Sox14 | TF | 169 ± 27 | 21 ± 21.8 | 1.11 ± 1.92 | 3.01 | 2.32E-03 | 7.24 | 1.52E-24 | nd |
| Tac1 | NP | 3017 ± 220 | 399 ± 92 | 848 ± 210 | 2.92 | 2.22E-48 | 1.82 | 1.09E-21 | |
| Trh | NP | 274 ± 44 | 37 ± 30 | 36.1 ± 16.6 | 2.88 | 1.43E-04 | 2.91 | 2.38E-14 | |

**Table S3.** Most enriched genes in vGAT vs. vGluT2. Group: TF = transcription factor, NP = neuropeptide, pNP = putative neuropeptide (classification of neuropeptides and putative neuropeptides is based on Burbach (2011)), NF = neurotrophic factor, NPR = neuropeptide receptor. sc data = single-cell data from Haring et al. (2018): nd = not detected, ng = not grouped.

|  |  | vGluT2<br>[norm. count] | vGAT<br>[norm. count] | Gad67<br>[norm. count] | vGluT2 vs. vGAT |  | vGluT2 vs. Gad67 |  |  |
| --- | --- | --- | --- | --- | --- | --- | --- | --- | --- |
| Gene | Group | Mean ± Stddev | Mean ± Stddev | Mean ± Stddev | log2 Ratio | FDR | log2 Ratio | FDR | sc Data |
| Tfap2b | TF | 30.6 ± 17.1 | 1455 ± 48 | 717 ± 60 | -5.58 | 2.29E-83 | -4.57 | 1.97E-48 |  |
| Pax8 | TF | 63.7 ± 9 | 2726 ± 338 | 3115 ± 419 | -5.43 | 1.32E-140 | -5.63 | 1.05E-171 |  |
| Sall3 | TF | 201 ± 63 | 5381 ± 754 | 6224 ± 804 | -4.75 | 3.76E-92 | -4.97 | 5.74E-122 |  |
| Slc32a1 |  | 734 ± 397 | 19147 ± 2462 | 20990 ± 2384 | -4.71 | 9.09E-39 | -4.85 | 1.12E-45 |  |
| Ighg3 |  | 0.7 ± 1.21 | 20 ± 3 | 124 ± 11.5 | -4.69 | 1.24E-01 | -7.39 | 8.67E-18 | nd |
| Gad2 |  | 802 ± 111 | 18347 ± 3188 | 26077 ± 1196 | -4.52 | 4.58E-91 | -5.04 | 5.70E-156 |  |
| Gad1 |  | 823 ± 140 | 18310 ± 4760 | 40767 ± 3643 | -4.48 | 1.69E-70 | -5.64 | 1.19E-127 |  |
| Pax2 | TF | 321 ± 168 | 6971 ± 499 | 7070 ± 544 | -4.45 | 3.30E-50 | -4.48 | 8.72E-56 |  |
| Pnoc | NP | 116 ± 30 | 2499 ± 184 | 3953 ± 558 | -4.44 | 7.27E-92 | -5.11 | 2.70E-129 |  |
| Lhx5 | TF | 59.4 ± 14.7 | 1129 ± 164 | 1129 ± 121 | -4.24 | 6.52E-59 | -4.24 | 3.88E-73 |  |
| Gata3 | TF | 6.8 ± 6.2 | 128 ± 45 | 18 ± 3.5 | -4.18 | 2.17E-06 | -1.39 | 6.05E-01 | ng |
| Slc6a5 |  | 1838 ± 823 | 32313 ± 2418 | 33567 ± 5237 | -4.14 | 3.30E-35 | -4.20 | 4.87E-43 |  |
| Gbx1 | TF | 67.3 ± 19.2 | 1172 ± 90 | 1359 ± 102 | -4.14 | 3.10E-58 | -4.36 | 6.33E-80 |  |
| Tal1 | TF | 11.8 ± 6.4 | 203.6 ± 11 | 21 ± 22.2 | -4.06 | 9.14E-15 | -0.83 | 7.89E-01 | ng |
| Dmkn |  | 5.6 ± 7.9 | 90.9 ± 12.5 | 123 ± 28 | -4.04 | 1.31E-03 | -4.49 | 1.13E-06 |  |
| Pax5 | TF | 13.4 ± 9.9 | 203 ± 66 | 204 ± 121 | -3.91 | 1.47E-10 | -3.92 | 4.29E-10 | nd |
| Aqp6 |  | 6.6 ± 7.4 | 78 ± 51 | 174 ± 32 | -3.57 | 2.34E-02 | -4.77 | 8.88E-12 |  |
| Dmrt3 | TF | 16.2 ± 17.6 | 189 ± 22.2 | 112 ± 29 | -3.57 | 7.82E-06 | -2.81 | 2.75E-03 | nd |
| Pkd1l2 |  | 13.5 ± 5.3 | 159 ± 27 | 0 ± 0 | -3.53 | 2.11E-09 | 6.84 | 1.80E-02 |  |
| Slc6a1 |  | 1513 ± 451 | 15617 ± 4052 | 26070 ± 3382 | -3.37 | 1.05E-37 | -4.12 | 2.78E-69 |  |
| Cdh3 |  | 64 ± 29 | 651 ± 253 | 1336 ± 184 | -3.36 | 4.75E-17 | -4.41 | 3.68E-58 |  |
| Grb7 |  | 7.3 ± 6.9 | 73 ± 36 | 84 ± 38 | -3.33 | 2.46E-02 | -3.54 | 1.31E-03 | ng |
| Gata2 | TF | 11.6 ± 9.5 | 109 ± 36 | 12.5 ± 10.3 | -3.23 | 3.46E-04 | -0.11 | 9.77E-01 | ng |
| Pkd2l1 |  | 41 ± 30 | 368 ± 110 | 30.4 ± 20.6 | -3.17 | 3.57E-11 | 0.42 | 8.20E-01 | ng |
| Alox12b |  | 15.2 ± 12.5 | 133 ± 72 | 202 ± 105 | -3.13 | 2.46E-04 | -3.73 | 8.08E-09 |  |
| Htr3a |  | 90.8 ± 21.7 | 752 ± 124 | 428 ± 75 | -3.05 | 1.11E-28 | -2.25 | 1.52E-15 |  |
| Asic4 |  | 63 ± 42 | 495 ± 31 | 610 ± 58 | -2.99 | 2.27E-13 | -3.30 | 1.25E-18 |  |
| Slc30a3 |  | 585 ± 201 | 4564 ± 487 | 4491 ± 486 | -2.97 | 6.37E-40 | -2.96 | 1.78E-44 |  |
| Ntsr1 | NPR | 63.3 ± 15.9 | 480 ± 34 | 726 ± 64 | -2.92 | 1.85E-22 | -3.53 | 1.27E-45 |  |
| Actr5 |  | 58.4 ± 21.8 | 439 ± 81 | 65.4 ± 19.3 | -2.90 | 1.60E-16 | -0.16 | 9.00E-01 |  |
| Gpc4 |  | 42.3 ± 17 | 311 ± 48 | 602 ± 150 | -2.89 | 3.30E-14 | -3.85 | 1.86E-32 |  |
| Shisa3 |  | 38.7 ± 22.9 | 273 ± 26 | 246 ± 64 | -2.84 | 1.66E-10 | -2.69 | 1.33E-09 | ng |
| Klhl14 |  | 207.1 ± 5.1 | 1475 ± 154 | 1121 ± 4.4 | -2.83 | 8.97E-48 | -2.45 | 1.58E-42 |  |
| Neurod2 | TF | 259.8 ± 8.5 | 1842 ± 146 | 1826 ± 92 | -2.83 | 2.00E-55 | -2.82 | 7.20E-70 |  |
| Krt25 |  | 59.2 ± 3.9 | 414 ± 98 | 650 ± 196 | -2.81 | 1.40E-18 | -3.47 | 2.26E-34 |  |
| Lhx1 | TF | 154 ± 54 | 1063 ± 169 | 1133 ± 106 | -2.79 | 8.26E-25 | -2.89 | 6.51E-33 |  |
| Neurod6 | TF | 141 ± 76 | 918 ± 132 | 659 ± 107 | -2.72 | 1.94E-15 | -2.24 | 3.56E-11 |  |
| C130060K24Rik |  | 14.5 ± 8.2 | 94 ± 67 | 93 ± 34 | -2.68 | 1.37E-02 | -2.67 | 7.61E-04 | ng |
| Neurod1 | TF | 63 ± 26 | 391 ± 72 | 379 ± 58 | -2.64 | 7.82E-13 | -2.60 | 7.23E-15 |  |
| Lgals9 |  | 219 ± 39 | 1343 ± 160 | 1813 ± 243 | -2.62 | 5.16E-34 | -3.06 | 3.38E-59 |  |
| Ghsr |  | 43.4 ± 22.2 | 263 ± 57 | 255 ± 48 | -2.62 | 5.63E-09 | -2.58 | 2.60E-10 | ng |
| Rasgrp1 |  | 237 ± 27 | 1256 ± 300 | 1337 ± 195 | -2.41 | 3.56E-25 | -2.50 | 9.85E-40 |  |
| Erich2 |  | 7 ± 6.8 | 37.3 ± 14.9 | 154 ± 30 | -2.39 | 3.52E-01 | -4.43 | 4.02E-11 | ng |
| Islr2 |  | 32.1 ± 3.1 | 163 ± 68 | 248 ± 33 | -2.35 | 4.53E-05 | -2.96 | 4.51E-18 |  |
| Fblim1 |  | 42.2 ± 19.1 | 214 ± 28 | 179 ± 35 | -2.35 | 2.52E-07 | -2.10 | 4.24E-06 | ng |
| Irx1 | TF | 23.7 ± 21.4 | 111.2 ± 20.4 | 216 ± 52 | -2.25 | 1.30E-01 | -3.21 | 2.28E-04 | ng |
| Nrl | TF | 28 ± 34 | 130 ± 31 | 127 ± 45 | -2.21 | 4.22E-02 | -2.18 | 2.00E-02 |  |
| Kcnf1 |  | 102.2 ± 24.4 | 466 ± 55 | 685 ± 156 | -2.19 | 1.02E-13 | -2.76 | 2.02E-25 |  |
| Gbx2 | TF | 88.7 ± 14.1 | 399 ± 119 | 348 ± 84 | -2.18 | 1.95E-10 | -1.99 | 2.18E-11 |  |
| Irx2 | TF | 23.1 ± 4.2 | 102 ± 50 | 244.3 ± 22 | -2.14 | 1.06E-02 | -3.40 | 3.21E-21 | ng |

**Table S4.**
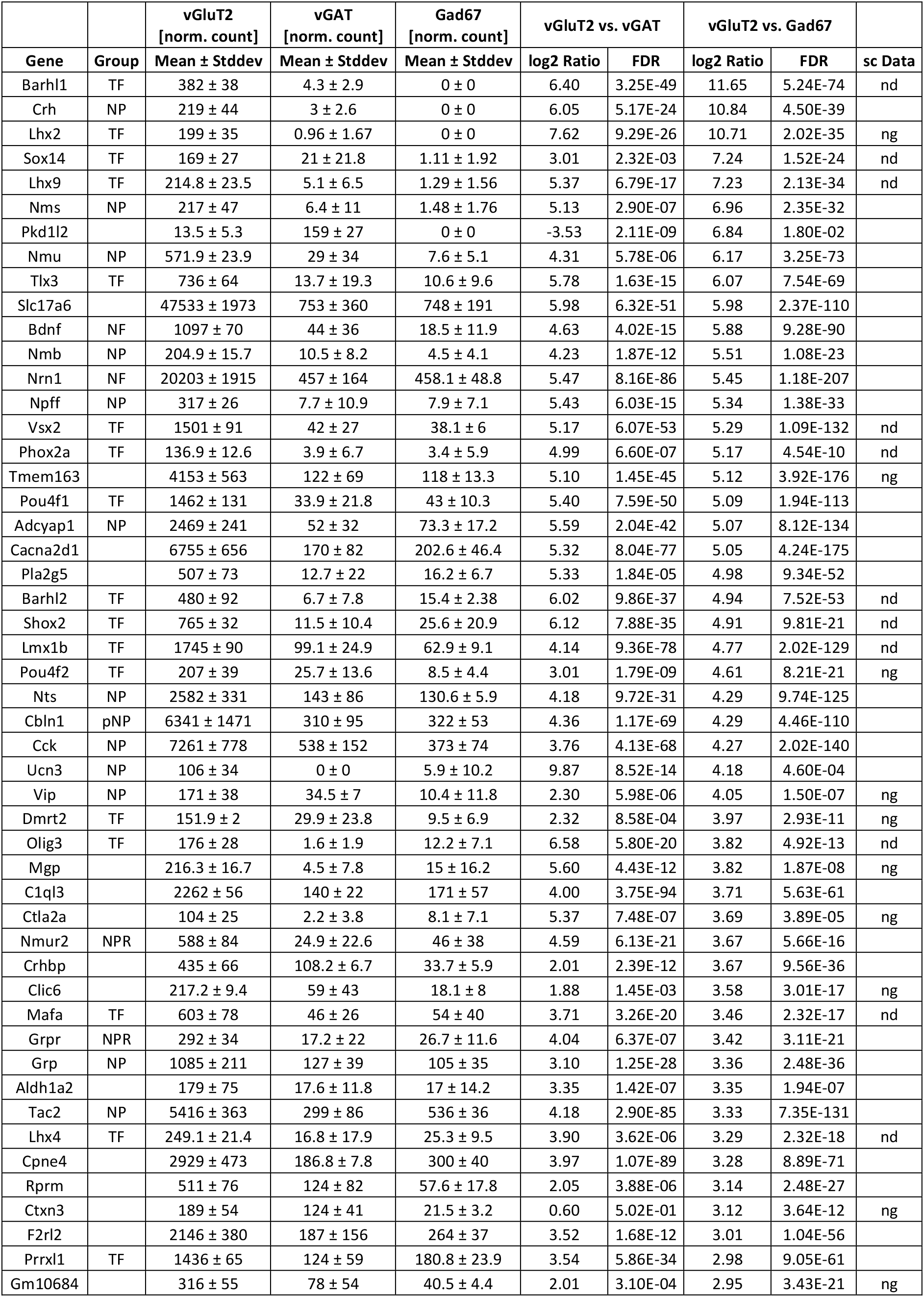
Most enriched genes in vGluT2 vs. Gad67. Group: TF = transcription factor, NP = neuropeptide, pNP = putative neuropeptide (classification of neuropeptides and putative neuropeptides is based on Burbach (2011)), NF = neurotrophic factor, NPR = neuropeptide receptor. sc data = single-cell data from Haring et al. (2018): nd = not detected, ng = not grouped.

**Table S5.** Most enriched genes in Gad67 vs. vGluT2. Group: TF = transcription factor, NP = neuropeptide, pNP = putative neuropeptide (classification of neuropeptides and putative neuropeptides is based on Burbach (2011)), NF = neurotrophic factor, NPR = neuropeptide receptor. sc data = single-cell data from Haring et al. (2018): nd = not detected, ng = not grouped.

|  |  | vGluT2<br>[norm. count] | vGAT<br>[norm. count] | Gad67<br>[norm. count] | vGluT2 vs. vGAT |  | vGluT2 vs. Gad67 |  |  |
| --- | --- | --- | --- | --- | --- | --- | --- | --- | --- |
| Gene | Group | Mean ± Stddev | Mean ± Stddev | Mean ± Stddev | log2 Ratio | FDR | log2 Ratio | FDR | sc Data |
| Ighg3 |  | 0.7 ± 1.21 | 20.0 ± 3.0 | 124 ± 11.5 | -4.69 | 1.24E-01 | -7.39 | 8.67E-18 | nd |
| Gad1 |  | 823 ± 140 | 18310 ± 4760 | 40767 ± 3643 | -4.48 | 1.69E-70 | -5.64 | 1.19E-127 |  |
| Pax8 | TF | 63.7 ± 9 | 2726 ± 338 | 3115 ± 419 | -5.43 | 1.32E-140 | -5.63 | 1.05E-171 |  |
| Chrna2 |  | 16.4 ± 14.6 | 9.28 ± 0.51 | 606 ± 100 | 0.80 | 9.30E-01 | -5.26 | 1.09E-16 | nd |
| Pnoc | NP | 116 ± 30 | 2499 ± 184 | 3953 ± 558 | -4.44 | 7.27E-92 | -5.11 | 2.70E-129 |  |
| Gad2 |  | 802 ± 111 | 18347 ± 3188 | 26077 ± 1196 | -4.52 | 4.58E-91 | -5.04 | 5.70E-156 |  |
| Sall3 | TF | 201 ± 63 | 5381 ± 754 | 6224 ± 804 | -4.75 | 3.76E-92 | -4.97 | 5.74E-122 |  |
| Slc32a1 |  | 734 ± 397 | 19147 ± 2462 | 20990 ± 2384 | -4.71 | 9.09E-39 | -4.85 | 1.12E-45 |  |
| Aqp6 |  | 6.6 ± 7.4 | 78 ± 51 | 174 ± 32 | -3.57 | 2.34E-02 | -4.77 | 8.88E-12 |  |
| Tfap2b | TF | 30.6 ± 17.1 | 1455 ± 48 | 717 ± 60 | -5.58 | 2.29E-83 | -4.57 | 1.97E-48 |  |
| Dmkn |  | 5.6 ± 7.9 | 90.9 ± 12.5 | 123 ± 28 | -4.04 | 1.31E-03 | -4.49 | 1.13E-06 |  |
| Pax2 | TF | 321 ± 168 | 6971 ± 499 | 7070 ± 544 | -4.45 | 3.30E-50 | -4.48 | 8.72E-56 |  |
| Erich2 |  | 7 ± 6.8 | 37.3 ± 14.9 | 154 ± 30 | -2.39 | 3.52E-01 | -4.43 | 4.02E-11 | ng |
| Cdh3 |  | 64 ± 29 | 651 ± 253 | 1336 ± 184 | -3.36 | 4.75E-17 | -4.41 | 3.68E-58 |  |
| Gbx1 | TF | 67.3 ± 19.2 | 1172 ± 90 | 1359 ± 102 | -4.14 | 3.10E-58 | -4.36 | 6.33E-80 |  |
| Lhx5 |  | 59.4 ± 14.7 | 1129 ± 164 | 1129 ± 121 | -4.24 | 6.52E-59 | -4.24 | 3.88E-73 |  |
| Slc6a5 |  | 1838 ± 823 | 32313 ± 2418 | 33567 ± 5237 | -4.14 | 3.30E-35 | -4.20 | 4.87E-43 |  |
| Slc6a1 |  | 1513 ± 451 | 15617 ± 4052 | 26070 ± 3382 | -3.37 | 1.05E-37 | -4.12 | 2.78E-69 | ng |
| Pax5 | TF | 13.4 ± 9.9 | 203 ± 66 | 204 ± 121 | -3.91 | 1.47E-10 | -3.92 | 4.29E-10 | nd |
| Gpc4 |  | 42.3 ± 17 | 311 ± 48 | 602 ± 150 | -2.89 | 3.30E-14 | -3.85 | 1.86E-32 |  |
| Alox12b |  | 15.2 ± 12.5 | 133 ± 72 | 202 ± 105 | -3.13 | 2.46E-04 | -3.73 | 8.08E-09 |  |
| Grb7 |  | 7.3 ± 6.9 | 73 ± 36 | 84 ± 38 | -3.33 | 2.46E-02 | -3.54 | 1.31E-03 | ng |
| Ntsr1 | NPR | 63.3 ± 15.9 | 480 ± 34 | 726 ± 64 | -2.92 | 1.85E-22 | -3.53 | 1.27E-45 |  |
| Krt25 |  | 59.2 ± 3.9 | 414 ± 98 | 650 ± 196 | -2.81 | 1.40E-18 | -3.47 | 2.26E-34 |  |
| Irx2 | TF | 23.1 ± 4.2 | 102 ± 50 | 244.3 ± 22 | -2.14 | 1.06E-02 | -3.40 | 3.21E-21 | ng |
| Slc12a7 |  | 21.2 ± 13.7 | 90 ± 50 | 215.3 ± 7.2 | -2.08 | 7.18E-02 | -3.35 | 4.56E-13 |  |
| Asic4 |  | 63 ± 42 | 495 ± 31 | 610 ± 58 | -2.99 | 2.27E-13 | -3.30 | 1.25E-18 |  |
| Irx1 | TF | 23.7 ± 21.4 | 111.2 ± 20.4 | 216 ± 52 | -2.25 | 1.30E-01 | -3.21 | 2.28E-04 | ng |
| Otp | TF | 112 ± 92 | 429 ± 96 | 967 ± 82 | -1.95 | 3.03E-04 | -3.13 | 6.67E-13 | nd |
| Lgals9 |  | 219 ± 39 | 1343 ± 160 | 1813 ± 243 | -2.62 | 5.16E-34 | -3.06 | 3.38E-59 |  |
| Adamtsl2 |  | 19.8 ± 2.7 | 58 ± 26 | 154 ± 58 | -1.56 | 3.22E-01 | -2.97 | 8.24E-09 |  |
| Islr2 |  | 32.1 ± 3.1 | 163 ± 68 | 248 ± 33 | -2.35 | 4.53E-05 | -2.96 | 4.51E-18 |  |
| Slc30a3 |  | 585 ± 201 | 4564 ± 487 | 4491 ± 486 | -2.97 | 6.37E-40 | -2.96 | 1.78E-44 |  |
| Crabp1 |  | 689 ± 32 | 2431 ± 336 | 5210 ± 543 | -1.82 | 4.04E-25 | -2.93 | 2.70E-101 |  |
| Lhx1 | TF | 154 ± 54 | 1063 ± 169 | 1133 ± 106 | -2.79 | 8.26E-25 | -2.89 | 6.51E-33 |  |
| Ppef1 |  | 19.2 ± 10.3 | 64.9 ± 8.7 | 140 ± 32 | -1.75 | 2.12E-01 | -2.86 | 1.11E-06 |  |
| Nfatc1 | TF | 32.1 ± 14.3 | 135 ± 67 | 222.9 ± 11 | -2.09 | 7.21E-03 | -2.83 | 8.07E-12 |  |
| Neurod2 | TF | 259.8 ± 8.5 | 1842 ± 146 | 1826 ± 92 | -2.83 | 2.00E-55 | -2.82 | 7.20E-70 |  |
| Dmrt3 | TF | 16.2 ± 17.6 | 189 ± 22.2 | 112 ± 29 | -3.57 | 7.82E-06 | -2.81 | 2.75E-03 | nd |
| Kcnf1 |  | 102.2 ± 24.4 | 466 ± 55 | 685 ± 156 | -2.19 | 1.02E-13 | -2.76 | 2.02E-25 |  |
| Shisa3 |  | 38.7 ± 22.9 | 273 ± 26 | 246 ± 64 | -2.84 | 1.66E-10 | -2.69 | 1.33E-09 | ng |
| C130060K24Rik |  | 14.5 ± 8.2 | 94 ± 67 | 93 ± 34 | -2.68 | 1.37E-02 | -2.67 | 7.61E-04 | ng |
| Neurod1 | TF | 63 ± 26 | 391 ± 72 | 379 ± 58 | -2.64 | 7.82E-13 | -2.60 | 7.23E-15 |  |
| En1 | TF | 47.1 ± 11.6 | 164 ± 30 | 281 ± 119 | -1.80 | 5.41E-04 | -2.59 | 2.92E-11 | nd |
| Ghsr |  | 43.4 ± 22.2 | 263 ± 57 | 255 ± 48 | -2.62 | 5.63E-09 | -2.58 | 2.60E-10 | ng |
| Rasgrp1 |  | 237 ± 27 | 1256 ± 300 | 1337 ± 195 | -2.41 | 3.56E-25 | -2.50 | 9.85E-40 |  |
| Irgm1 |  | 127.8 ± 12.4 | 559 ± 115 | 715 ± 124 | -2.13 | 2.50E-14 | -2.49 | 5.19E-27 |  |
| Klhl14 |  | 207.1 ± 5.1 | 1475 ± 154 | 1121 ± 4.4 | -2.83 | 8.97E-48 | -2.45 | 1.58E-42 |  |
| Pdyn | NP | 384 ± 77 | 966 ± 280 | 2075 ± 385 | -1.33 | 1.94E-06 | -2.44 | 1.46E-33 |  |
| Npy | NP | 1262 ± 325 | 3605 ± 812 | 6622 ± 880 | -1.52 | 1.24E-10 | -2.41 | 3.66E-38 |  |

## Neurotransmitter receptor expression

### Glutamate receptors

**Table S5.** Ionotropic glutamate receptors (AMPA, kainate and NMDA receptors). Significant (FDR ≤ 0.05) and | log2 ratio | ≥ 1 enrichment values are highlighted in **bold**.

|  | vGluT2<br>[norm. count] | vGAT<br>[norm. count] | Gad67<br>[norm. count] | vGluT2 vs. vGAT |  | vGluT2 vs. Gad67 |  |
| --- | --- | --- | --- | --- | --- | --- | --- |
| Gene | Mean ± Stddev | Mean ± Stddev | Mean ± Stddev | log2<br>Ratio | FDR | log2<br>Ratio | FDR |
| Gria1 | 3853 ± 429 | 6189 ± 1252 | 9054 ± 821 | -0.69 | **1.92E-03** | **-1.24** | **4.49E-19** |
| Gria2 | 7524 ± 295 | 5010 ± 579 | 6896 ± 1352 | 0.59 | **2.98E-03** | 0.11 | 7.16E-01 |
| Gria3 | 3327 ± 203 | 3423 ± 208 | 3229 ± 304 | -0.04 | 9.66E-01 | 0.03 | 9.17E-01 |
| Gria4 | 4879 ± 727 | 4476 ± 238 | 4437 ± 544 | 0.12 | 8.68E-01 | 0.13 | 6.66E-01 |
| Grik1 | 200 ± 56 | 465 ± 183 | 453 ± 94 | **-1.22** | **2.32E-03** | **-1.19** | **8.15E-05** |
| Grik2 | 1074 ± 299 | 2411 ± 362 | 2676 ± 353 | **-1.17** | **3.08E-07** | **-1.33** | **7.94E-11** |
| Grik3 | 737 ± 97 | 2753 ± 114 | 3288 ± 275 | **-1.90** | **9.00E-30** | **-2.17** | **2.74E-44** |
| Grik4 | 651 ± 67 | 721 ± 179 | 735 ± 44 | -0.15 | 8.97E-01 | -0.19 | 5.66E-01 |
| Grik5 | 2341 ± 392 | 2481 ± 229 | 2289 ± 396 | -0.09 | 9.28E-01 | 0.02 | 9.66E-01 |
| Grin1 | 16293 ± 2625 | 14053 ± 2385 | 17130 ± 3003 | 0.21 | 8.29E-01 | -0.08 | 8.54E-01 |
| Grin2a | 3811 ± 484 | 3237 ± 352 | 3463 ± 703 | 0.23 | 5.72E-01 | 0.13 | 7.07E-01 |
| Grin2b | 4048 ± 192 | 3630 ± 254 | 4605 ± 535 | 0.16 | 7.71E-01 | -0.20 | 3.65E-01 |
| Grin2c | 41.4 ± 18.9 | 97.4 ± 21.4 | 116.4 ± 7.8 | **-1.24** | 2.43E-01 | **-1.51** | **1.25E-02** |
| Grin2d | 1775 ± 178 | 2062 ± 121 | 2485 ± 454 | -0.22 | 6.25E-01 | -0.50 | **1.75E-02** |
| Grin3a | 3791 ± 271 | 1965 ± 135 | 2127 ± 174 | 0.95 | **1.79E-09** | 0.82 | **2.86E-09** |
| Grin3b | 47.5 ± 11.5 | 49.0 ± 33.0 | 58.9 ± 26.6 | -0.06 | 9.92E-01 | -0.32 | 8.08E-01 |

**Table S6.** Metabotropic glutamate receptors. Significant (FDR ≤ 0.05) log2 ratio ? 1 enrichment values are highlighted in **bold**. NA = not applicable.

|  | vGluT2<br>[norm. count] | vGAT<br>[norm. count] | Gad67<br>[norm. count] | vGluT2 vs. vGAT |  | vGluT2 vs. Gad67 |  |
| --- | --- | --- | --- | --- | --- | --- | --- |
| Gene | Mean ± Stddev | Mean ± Stddev | Mean ± Stddev | log2<br>Ratio | FDR | log2<br>Ratio | FDR |
| Grm1 | 2407 ± 156 | 1632 ± 116 | 1912 ± 419 | 0.56 | **5.76E-03** | 0.32 | 2.15E-01 |
| Grm2 | 157.4 ± 23.5 | 184 ± 109 | 315.9 ± 8.2 | -0.22 | 9.27E-01 | **-1.02** | **2.70E-04** |
| Grm3 | 424 ± 79 | 946 ± 237 | 1243 ± 198 | **-1.16** | **1.86E-05** | **-1.57** | **1.06E-14** |
| Grm4 | 2116 ± 315 | 1574 ± 110 | 1714 ± 263 | 0.43 | 1.03E-01 | 0.29 | 2.55E-01 |
| Grm5 | 5322 ± 309 | 7181 ± 1239 | 9795 ± 1607 | -0.43 | 1.07E-01 | -0.89 | **2.53E-08** |
| Grm6 | 0 ± 0 | 0 ± 0 | 0 ± 0 | 0.00 | NA | 0.00 | NA |
| Grm7 | 1559 ± 184 | 1732 ± 161 | 1985 ± 194 | -0.15 | 8.25E-01 | -0.36 | 9.15E-02 |
| Grm8 | 420 ± 56 | 447 ± 29 | 549 ± 176 | -0.09 | 9.48E-01 | -0.40 | 3.07E-01 |

### GABA receptors

**Table S7.** Ionotropic GABA receptors (GABA_A_ and −ρ receptor). Significant (FDR ≤ 0.05) and | log2 ratio| ≥ 1 enrichment values are highlighted in **bold**. NA = not applicable.

|  | vGluT2<br>[norm. count] | vGAT<br>[norm. count] | Gad67<br>[norm. count] | vGluT2 vs. vGAT |  | vGluT2 vs. Gad67 |  |
| --- | --- | --- | --- | --- | --- | --- | --- |
| Gene | Mean ± Stddev | Mean ± Stddev | Mean ± Stddev | log2<br>Ratio | FDR | log2<br>Ratio | FDR |
| Gabra1 | 4236 ± 493 | 4361 ± 357 | 5504 ± 592 | -0.04 | 9.68E-01 | -0.39 | **2.87E-02** |
| Gabra2 | 977 ± 83 | 643 ± 93 | 638.4 ± 19.4 | 0.60 | **2.39E-02** | 0.60 | **1.76E-03** |
| Gabra3 | 5357 ± 498 | 4949 ± 585 | 6146 ± 837 | 0.11 | 8.84E-01 | -0.21 | 3.57E-01 |
| Gabra4 | 653 ± 39 | 555 ± 63 | 530.4 ± 22.5 | 0.23 | 7.36E-01 | 0.29 | 3.08E-01 |
| Gabra5 | 5260 ± 959 | 2574 ± 84 | 2355 ± 322 | **1.03** | **2.11E-09** | **1.15** | **3.97E-12** |
| Gabra6 | 0 ± 0 | 0 ± 0 | 0 ± 0 | 0.00 | NA | 0.00 | NA |
| Gabrb1 | 2706 ± 238 | 2797 ± 353 | 3234 ± 418 | -0.05 | 9.64E-01 | -0.27 | 2.01E-01 |
| Gabrb2 | 1637 ± 81 | 1152 ± 94 | 1364 ± 183 | 0.51 | **2.12E-02** | 0.25 | 3.06E-01 |
| Gabrb3 | 11663 ± 1308 | 9704 ± 1598 | 10398 ± 1382 | 0.26 | 6.50E-01 | 0.15 | 6.18E-01 |
| Gabrg1 | 593.2 ± 16.6 | 511 ± 119 | 941 ± 211 | 0.21 | 8.11E-01 | -0.68 | **4.50E-03** |
| Gabrg2 | 6631 ± 484 | 5536 ± 958 | 7043 ± 1243 | 0.26 | 5.52E-01 | -0.10 | 7.60E-01 |
| Gabrg3 | 1553 ± 158 | 1431 ± 452 | 1736 ± 237 | 0.12 | 9.24E-01 | -0.17 | 5.58E-01 |
| Gabrd | 36.2 ± 11.0 | 40 ± 52 | 28.8 ± 6.0 | -0.13 | 9.89E-01 | 0.32 | 8.51E-01 |
| Gabre | 97.6 ± 7.8 | 44.1 ± 11.7 | 58.7 ± 24.2 | **1.14** | 2.46E-01 | 0.72 | 3.44E-01 |
| Gabrp | 0 ± 0 | 0 ± 0 | 4.1 ± 7.0 | 0.00 | NA | **-5.13** | NA |
| Gabrq | 538.8 ± 24.7 | 217 ± 71 | 330 ± 85 | **1.31** | **2.06E-05** | 0.69 | **1.65E-02** |
| Gabrr1 | 12.1 ± 11.5 | 0.32 ± 0.56 | 0 ± 0 | **4.85** | 3.62E-01 | **6.68** | 6.79E-02 |
| Gabrr2 | 8.1 ± 8.3 | 17.6 ± 15.3 | 8.3 ± 9.1 | **-1.13** | 9.06E-01 | -0.06 | NA |
| Gabrr3 | 0 ± 0 | 0 ± 0 | 0 ± 0 | 0.00 | NA | 0.00 | NA |

**Table S8.** Metabotropic GABA receptors (GABAB). Significant (FDR ≤ 0.05) and |log2 ratio| ? 1 enrichment values are highlighted in **bold**.

|  | vGluT2<br>[norm. count] | vGAT<br>[norm. count] | Gad67<br>[norm. count] | vGluT2 vs. vGAT |  | vGluT2 vs. Gad67 |  |
| --- | --- | --- | --- | --- | --- | --- | --- |
| Gene | Mean ± Stddev | Mean ± Stddev | Mean ± Stddev | log2<br>Ratio | FDR | log2<br>Ratio | FDR |
| Gabbr1 | 26433 ± 4708 | 26183 ± 3572 | 28147 ± 4127 | 0.01 | 9.96E-01 | -0.10 | 8.39E-01 |
| Gabbr2 | 10007 ± 784 | 9451 ± 171 | 10552 ± 1623 | 0.08 | 9.36E-01 | -0.09 | 8.01E-01 |

### Glycine receptors

**Table S9.**
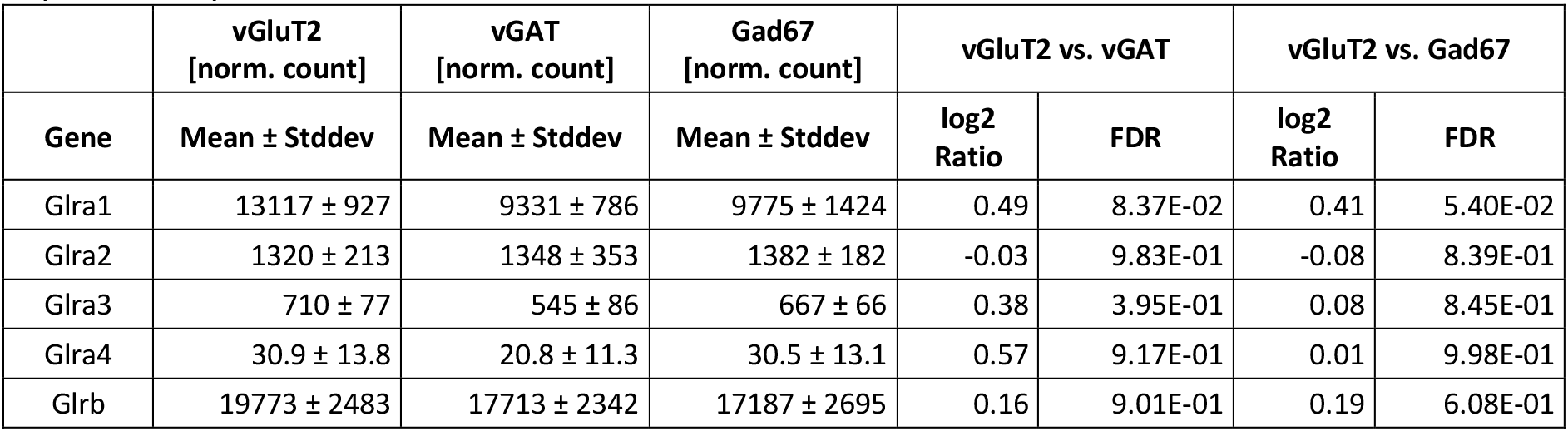
Glycine receptors. Significant (FDR ≤ 0.05) and |log2 ratio| ? 1 enrichment values are highlighted in **bold**.

### Acetylcholine receptors

**Table S10.** Ionotropic acetylcholine receptors (nicotinic). Significant (FDR ≤ 0.05) and |log2 ratio| ≥ 1 enrichment values are highlighted in **bold**. NA = not applicable.

|  | vGluT2<br>[norm. count] | vGAT<br>[norm. count] | Gad67<br>[norm. count] | vGluT2 vs. vGAT |  | vGluT2 vs. Gad67 |  |
| --- | --- | --- | --- | --- | --- | --- | --- |
| Gene | Mean ± Stddev | Mean ± Stddev | Mean ± Stddev | log2<br>Ratio | FDR | log2<br>Ratio | FDR |
| Chrna1 | 4.5 ± 7.8 | 3.8 ± 6.7 | 0 ± 0 | 0.22 | NA | **5.27** | NA |
| Chrna2 | 16.4 ± 14.6 | 9.3 ± 0.5 | 606 ± 100 | 0.80 | 9.30E-01 | **-5.26** | **1.09E-16** |
| Chrna3 | 117 ± 39 | 211 ± 53 | 420.9 ± 15.4 | -0.86 | 1.71E-01 | **-1.87** | **4.08E-10** |
| Chrna4 | 4316 ± 654 | 3537 ± 467 | 3457 ± 643 | 0.29 | 4.33E-01 | 0.31 | 1.95E-01 |
| Chrna5 | 26.2 ± 16.7 | 8.9 ± 8.5 | 97.6 ± 22.5 | **1.56** | 7.82E-01 | **-1.92** | **1.79E-02** |
| Chrna6 | 89.1 ± 43.0 | 3.3 ± 2.9 | 0 ± 0 | **4.66** | **1.59E-05** | **9.54** | **1.05E-12** |
| Chrna7 | 513 ± 114 | 493 ± 72 | 674 ± 239 | 0.05 | 9.75E-01 | -0.41 | 3.58E-01 |
| Chrna9 | 0 ± 0 | 0 ± 0 | 3.0 ± 5.2 | 0.00 | NA | **-4.73** | NA |
| Chrna10 | 0 ± 0 | 3.9 ± 6.7 | 6.6 ± 6.5 | **-5.13** | NA | **-5.82** | NA |
| Chrn b1 | 4.3 ± 7.4 | 0 ± 0 | 0 ± 0 | **5.28** | NA | **5.21** | NA |
| Chrn b2 | 1921 ± 238 | 1820 ± 286 | 2018 ± 214 | 0.08 | 9.43E-01 | -0.08 | 8.08E-01 |
| Chrn b3 | 38.1 ± 14.1 | 14.0 ± 11.3 | 8.5 ± 7.9 | **1.42** | 6.81E-01 | **2.15** | 2.25E-01 |
| Chrn b4 | 11.0 ± 3.7 | 58 ± 28 | 95.5 ± 21.1 | **-2.39** | 8.09E-02 | **-3.14** | **8.46E-06** |
| Chrnd | 0 ± 0 | 0 ± 0 | 0 ± 0 | 0.00 | NA | 0.00 | NA |
| Chrne | 0 ± 0 | 0 ± 0 | 0 ± 0 | 0.00 | NA | 0.00 | NA |
| Chrng | 3.8 ± 6.6 | 0 ± 0 | 0 ± 0 | **5.10** | NA | **5.03** | NA |

**Table S11.** Metabotropic acetylcholine receptors (muscarinic). Significant (FDR ≤ 0.05) and |log2 ratio| ≥ 1 enrichment values are highlighted in **bold**. NA = not applicable.

|  | vGluT2<br>[norm. count] | vGAT<br>[norm. count] | Gad67<br>[norm. count] | vGluT2 vs. vGAT |  | vGluT2 vs. Gad67 |  |
| --- | --- | --- | --- | --- | --- | --- | --- |
| Gene | Mean ± Stddev | Mean ± Stddev | Mean ± Stddev | log2<br>Ratio | FDR | log2<br>Ratio | FDR |
| Chrm1 | 25.6 ± 7.2 | 30.2 ± 6.1 | 19.3 ± 7.0 | -0.24 | 9.70E-01 | 0.39 | 8.53E-01 |
| Chrm2 | 5176 ± 811 | 4818 ± 667 | 5664 ± 782 | 0.10 | 9.13E-01 | -0.14 | 6.30E-01 |
| Chrm3 | 497 ± 148 | 1026 ± 160 | 1183 ± 141 | **-1.05** | **2.63E-04** | **-1.27** | **1.17E-07** |
| Chrm4 | 89 ± 34 | 56 ± 28 | 130.5 ± 19.8 | 0.66 | 7.68E-01 | -0.57 | 3.88E-01 |
| Chrm5 | 177 ± 28 | 8.8 ± 8.4 | 23.1 ± 13.1 | **4.27** | **3.14E-09** | **2.91** | **7.15E-09** |

### Serotonin receptors

**Table S12.** Serotonin receptors. Significant (FDR ≤ 0.05) and |log2 ratio| ? 1 enrichment values are highlighted in **bold**. NA = not applicable.

|  | vGluT2<br>[norm. count] | vGAT<br>[norm. count] | Gad67<br>[norm. count] | vGluT2 vs. vGAT |  | vGluT2 vs. Gad67 |  |
| --- | --- | --- | --- | --- | --- | --- | --- |
| Gene | Mean ± Stddev | Mean ± Stddev | Mean ± Stddev | log2<br>Ratio | FDR | log2<br>Ratio | FDR |
| Htr1a | 598 ± 103 | 1503 ± 240 | 1627 ± 154 | **-1.33** | **2.38E-10** | **-1.46** | **6.27E-16** |
| Htr1b | 803 ± 175 | 529 ± 132 | 543 ± 75 | 0.60 | 1.23E-01 | 0.55 | **4.26E-02** |
| Htr1d | 102 ± 37 | 214 ± 46 | 468 ± 102 | **-1.08** | **4.51E-02** | **-2.22** | **9.50E-12** |
| Htr1f | 68 ± 28 | 115 ± 91 | 60.3 ± 8.1 | -0.75 | 7.50E-01 | 0.16 | 8.99E-01 |
| Htr2a | 315 ± 56 | 466 ± 115 | 510 ± 71 | -0.57 | 2.29E-01 | -0.71 | **1.03E-02** |
| Htr2b | 0 ± 0 | 0 ± 0 | 0 ± 0 | 0.00 | NA | 0.00 | NA |
| Htr2c | 6685 ± 1060 | 10081 ± 1679 | 10828 ± 846 | -0.59 | **2.50E-02** | -0.71 | **3.75E-05** |
| Htr3a | 90.8 ± 21.7 | 752 ± 124 | 428 ± 75 | **-3.05** | **1.11E-28** | **-2.25** | **1.52E-15** |
| Htr3b | 3.1 ± 5.4 | 7.7 ± 5.9 | 7.0 ± 6.7 | **-1.29** | NA | **-1.15** | NA |
| Htr4 | 197.5 ± 23.6 | 130.5 ± 23.8 | 167 ± 60 | 0.60 | 3.60E-01 | 0.23 | 7.16E-01 |
| Htr5a | 525 ± 41 | 784 ± 59 | 699 ± 78 | -0.58 | **3.30E-02** | -0.43 | 9.91E-02 |
| Htr5b | 12.7 ± 10.1 | 17.2 ± 15.6 | 23.7 ± 19.8 | -0.45 | 9.68E-01 | -0.92 | 7.10E-01 |
| Htr6 | 87.4 ± 18.1 | 66 ± 35 | 109 ± 35 | 0.39 | 8.88E-01 | -0.33 | 6.90E-01 |
| Htr7 | 804 ± 48 | 1069 ± 120 | 828 ± 85 | -0.41 | 1.77E-01 | -0.05 | 8.90E-01 |

### Adrenergic receptors

**Table S13.** Adrenergic receptors. Significant (FDR ≤ 0.05) and | log2 ratio| ≥ 1 enrichment values are highlighted in **bold**. NA = not applicable.

|  | vGluT2<br>[norm. count] | vGAT<br>[norm. count] | Gad67<br>[norm. count] | vGluT2 vs. vGAT |  | vGluT2 vs. Gad67 |  |
| --- | --- | --- | --- | --- | --- | --- | --- |
| Gene | Mean ± Stddev | Mean ± Stddev | Mean ± Stddev | log2<br>Ratio | FDR | log2<br>Ratio | FDR |
| Adra1a | 576 ± 61 | 503 ± 112 | 595 ± 153 | 0.19 | 8.52E-01 | -0.06 | 9.14E-01 |
| Adra1b | 498 ± 152 | 753 ± 98 | 743 ± 95 | -0.60 | 1.19E-01 | -0.59 | 5.25E-02 |
| Adra1d | 101 ± 28 | 110.0 ± 20.3 | 117 ± 49 | -0.13 | 9.66E-01 | -0.22 | 8.07E-01 |
| Adra2a | 1992 ± 386 | 942 ± 155 | 981 ± 189 | **1.08** | **6.89E-07** | **1.01** | **1.27E-06** |
| Adra2b | 1.40 ± 2.42 | 6.0 ± 10.5 | 26.8 ± 15.0 | **-2.05** | NA | **-4.21** | 5.84E-02 |
| Adra2c | 389 ± 103 | 329.5 ± 17.7 | 302 ± 82 | 0.24 | 8.29E-01 | 0.35 | 4.56E-01 |
| Adrb1 | 105 ± 46 | 204 ± 46 | 162 ± 84 | -0.97 | 1.02E-01 | -0.65 | 3.46E-01 |
| Adrb2 | 9.0 ± 11.4 | 3.8 ± 6.7 | 0 ± 0 | **1.21** | NA | **6.25** | NA |
| Adrb3 | 12.5 ± 12.5 | 30.8 ± 22.7 | 24.5 ± 3.2 | **-1.30** | 8.12E-01 | -0.99 | 7.08E-01 |

### Dopamine receptors

**Table S14.**
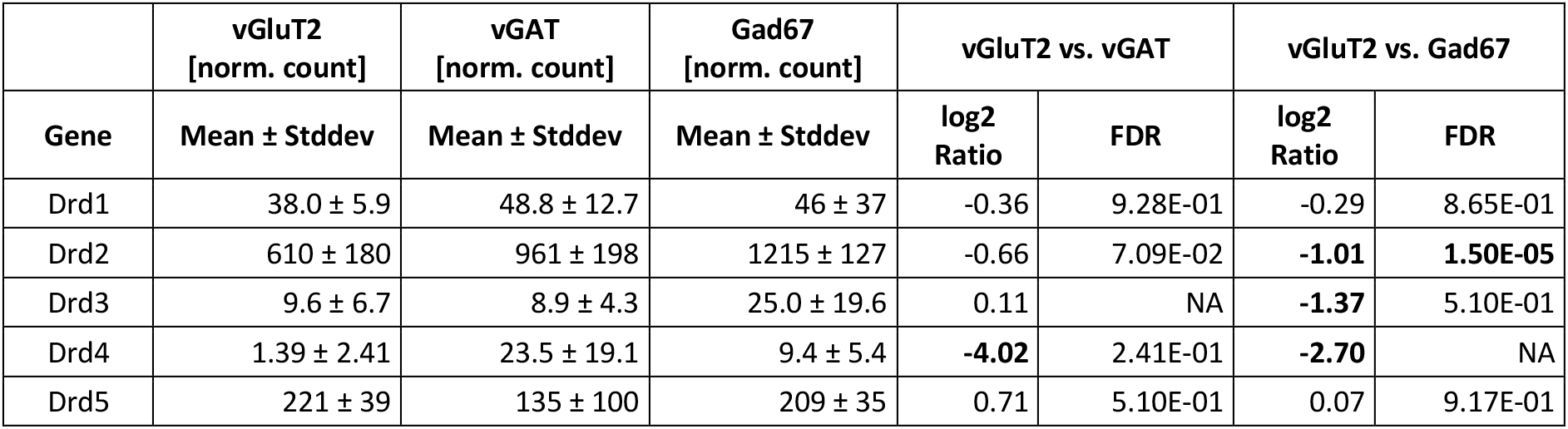
Dopamine receptors. Significant (FDR ≤ 0.05) and |log2 ratio| ≥ 1 enrichment values are highlighted in **bold**. NA = not applicable.

### Histamine receptors

**Table S15.** Histamine receptors. Significant (FDR ≤ 0.05) and |log2 ratio| ≥ 1 enrichment values are highlighted in **bold**. NA = not applicable.

|  | vGluT2<br>[norm. count] | vGAT<br>[norm. count] | Gad67<br>[norm. count] | vGluT2 vs. vGAT |  | vGluT2 vs. Gad67 |  |
| --- | --- | --- | --- | --- | --- | --- | --- |
| Gene | Mean ± Stddev | Mean ± Stddev | Mean ± Stddev | log2<br>Ratio | FDR | log2<br>Ratio | FDR |
| Hrh1 | 39.1 ± 9.8 | 35.2 ± 21.0 | 73 ± 38 | 0.14 | 9.81E-01 | -0.92 | 3.32E-01 |
| Hrh2 | 80.1 ± 21.4 | 77 ± 36 | 68.8 ± 7.8 | 0.05 | 9.92E-01 | 0.21 | 8.37E-01 |
| Hrh3 | 942 ± 181 | 1428 ± 79 | 1523 ± 94 | -0.60 | **1.36E-02** | -0.71 | **4.04E-04** |
| Hrh4 | 0 ± 0 | 0 ± 0 | 0 ± 0 | 0.00 | NA | 0.00 | NA |

